# Dynamic representations of faces in the human ventral visual stream link visual features to behaviour

**DOI:** 10.1101/394916

**Authors:** Diana C. Dima, Krish D. Singh

## Abstract

Humans can rapidly extract information from faces even in challenging viewing conditions, yet the neural representations supporting this ability are still not well understood. Here, we manipulated the presentation duration of backward-masked facial expressions and used magnetoencephalography (MEG) to investigate the computations underpinning rapid face processing. Multivariate analyses revealed two stages in face perception, with the ventral visual stream encoding facial features prior to facial configuration. When presentation time was reduced, the emergence of sustained featural and configural representations was delayed. Importantly, these representations explained behaviour during an expression recognition task. Together, these results describe the adaptable system linking visual features, brain and behaviour during face perception.

## Introduction

Our highly specialized face processing abilities are thought to be supported by feature-based face detection followed by configural processing (Calder, Young, Keane, & Dean, 2000; Maurer, Grand, & Mondloch, 2002). However, it is still unclear how the brain efficiently represents a high-dimensional array of relevant facial features, and how this helps accomplish a wide range of behavioural goals.

Although there is disagreement on the exact sequence of processing stages, face perception is generally thought to progress from isolated features to first-order configuration (the feature positioning common across all faces) and second-order configuration (the identity-specific spacing between features), with holistic processing linking these into a gestalt (Farah, Wilson, & Tanaka, 1998; Harris & Aguirre, 2008; Piepers & Robbins, 2012). On the other hand, some behavioural goals, such as identity recognition, may rely on facial features and not on holistic perception (Visconti Di Oleggio Castello, Wheeler, Cipolli, & Gobbini, 2017).

Furthermore, although the neural correlates of face perception have been reliably mapped in space and time, there is little agreement on how, where, and when specific computations are implemented. Both modular and distributed neural codes are thought to support face perception, with different computations being implemented within each of the ventral face-responsive areas (Grill-Spector, Weiner, Gomez, Stigliani, & Natu, 2018; Freiwald, Duchaine, & Yovel, 2016). For efficient information extraction, faces may be represented along low-dimensional axes based on features or topology (Henriksson, Mur, & Kriegeskorte, 2015; Leopold, O’Toole, Vetter, & Blanz, 2001); for example, a sparse identity code has been shown to predict neural responses to faces in primates (Chang & Tsao, 2017). However, it remains an open question how such codes adapt to task requirements and viewing conditions, and the dynamics of face feature representations are not well understood.

Here, we focused on the temporal dynamics of face representations during a challenging expression discrimination task, by combining magnetoencephalography (MEG) with multivariate pattern analyses and a rapid presentation paradigm. We manipulated the presentation duration of backward-masked faces, some of which were shown outside awareness, to disentangle face detection from expression processing. This allowed us to evaluate the impact of limiting visual input on representational dynamics, while keeping task demands constant. We used source-space representational similarity analysis (RSA) and variance partitioning to evaluate the contribution of visual features to MEG responses and behaviour.

We found that among the visual features tested, facial features and configuration were most strongly represented in the ventral stream and contributed to behaviour. The temporal dynamics of these representations changed in response to stimulus duration, suggesting that it is important to study visual feature coding in dynamic contexts and with high temporal resolution. Finally, despite a behavioural effect, a neural response to faces outside of awareness did not encode any of the stimulus features tested, highlighting the qualitative distinction between face detection and face categorization.

## Methods

### Participants

The participants were 25 healthy volunteers (16 female, age range 19-42, mean age 25.6 ± 5.39). All volunteers gave written consent to participate in the study in accordance with The Code of Ethics of the World Medical Association (Declaration of Helsinki). All procedures were approved by the ethics committee of the School of Psychology, Cardiff University.

### Stimuli

The stimulus set consisted of 20 faces with angry, neutral and happy expressions (10 female faces; model numbers: 2, 6, 7, 8, 9, 11, 14, 16, 17, 18, 22, 23, 25, 31, 34, 35, 36, 38, 39, 40) from the NIMSTIM database (Tottenham et al., 2009). The eyes were aligned using automated eye detection as implemented in the Matlab Computer Vision System toolbox (Mathworks, Inc., Natick, Massachusetts). An oval mask was used to crop the faces to a size of 378 × 252 pixels subtending 2.6 × 3.9 degrees of visual angle. All images were converted to grayscale. Their spatial frequency was matched by specifying the rotational average of the Fourier amplitude spectra as implemented in the SHINE toolbox (Willenbockel et al., 2010), and Fourier amplitude spectra for all faces were set to the average across the face set.

Masks and control stimuli were created by scrambling the phase of all face images in the Fourier domain. This was achieved by replacing the phase information in each of the images with phase information from a white noise image of equal size (Perry & Singh, 2014). To ensure matched low-level properties between face and control stimuli, pixel intensities were normalized between each image and its scrambled counterpart, using the minimum and maximum pixel intensity of the scrambled image.

### Experimental design

At the start of each trial, a white fixation cross was centrally presented on an isoluminant gray background. Its duration was pseudorandomly chosen from a uniform distribution between 1.3 and 1.6 s. A face stimulus was then centrally presented with a duration of either 10 ms, 30 ms or 150 ms; the stimulus was followed by a phase-scrambled mask with a duration of 190 ms, 170 ms or 50 ms respectively (for a constant total stimulus duration of 200 ms). In each block, 10 trials contained no face; instead, a phase-scrambled control stimulus was flashed for 10 ms and followed by another mask.

After a 500 ms delay intended to dissociate face perception from response preparation, participants had to correctly select the expression they had perceived out of three alternatives presented on screen (Figure 3A). They had 1.5 seconds to make a button press; if they were sure that no face had been presented, they could refrain from responding. The mapping of the response buttons to emotional expressions changed halfway through the experiment so as to ensure that emotional expression processing would not be confounded by specific motor preparation effects.

Next, participants had to rate how clearly they had seen the face using a 3-point scale starting from 0. They were instructed to only select 0 if no face had been perceived, 1 if they had perceived a face but not clearly, and 2 if they had clearly perceived the face. They had 2 seconds to make this response. Note that since the expression discrimination task was not forced-choice, references to awareness in this paper refer exclusively to subjective awareness, as indicated by perceptual ratings.

In each of four blocks, each face was presented once with each of the three possible stimulus durations. We thus collected 80 trials per condition, except for the control condition (containing no face) which only had 40 trials.

### Data acquisition

All participants with one exception acquired a whole-head structural MRI using a 1 mm isotropic Fast Spoiled Gradient-Recalled-Echo pulse sequence.

Whole-head MEG recordings were made using a 275-channel CTF radial gradiometer system (CTF, Vancouver, Canada) at a sampling rate of 1200 Hz. Four of the sensors were turned off due to excessive sensor noise. An additional 29 reference channels were recorded for noise rejection purposes and the primary sensors were analyzed as synthetic third-order gradiometers (Vrba & Robinson, 2001).

Stimuli were presented using a ProPixx projector system (VPixx Technologies, Saint-Bruno, Canada) with a refresh rate set to 100 Hz. Images were projected to a screen with a resolution of 1920 × 1080 pixels situated at a distance of 1.2 m from the participant. Recordings were made in four blocks of approximately 15 minutes each, separated by short breaks. The data were collected in 2.5 s epochs beginning 1 s prior to stimulus onset.

Participants were seated upright while viewing the stimuli and electromagnetic coils were attached to the nasion and pre-auricular points on the scalp in order to continuously monitor head position relative to a fixed coordinate system on the dewar. To help co-register the MEG data with the participants’ structural MRI scans, we defined the head shape of each subject using an ANT Xensor digitizer (ANT Neuro, Enschede, Netherlands). An Eyelink 1000 eye-tracker system (SR Research, Ottawa, Canada) with a sampling rate of 1000 Hz was used to track the subjects’ right pupil and corneal reflex.

### Behavioural analysis

In order to assess the effects of stimulus duration and face expression on behaviour, we calculated confusion matrices based on expression discrimination responses to each stimulus category (Figure 3). Performance was quantified as proportion correct trials after excluding trials with no response, and a rationalized arcsine transformation was applied prior to statistical analysis (Studebaker, 1985). We then performed a 3 × 3 repeated-measures ANOVA with factors *Duration* (levels: 10 ms, 30 ms, 150 ms) and *Expression* (levels: angry, happy, neutral).

### MEG multivariate pattern analysis (MVPA)

To test for differences between conditions present in multivariate patterns, we used a linear Support Vector Machine (SVM) classifier with L2 regularization and a box constraint *c* = 1. The classifier was implemented in Matlab using LibLinear (Fan, Chang, Hsieh, Wang, & Lin, 2008) and the Statistics and Machine Learning Toolbox (Mathworks, Inc.). We performed binary classification on (1) responses to neutral faces versus scrambled stimuli (face decoding); (2) all three pairs of emotional expressions (expression decoding).

For face decoding, time-resolved classification was performed separately for each stimulus duration. To assess the presence of subjectively non-conscious responses, the classification of faces presented for 10 ms was performed after excluding any trials reported as containing a face. To ensure that decoding results were not biased by stimulus repetitions or recognition of face identities across the training and test sets, cross-exemplar five-fold cross-validation was used to assess classification performance: the classifier was trained on 16 of the 20 face identities and 8 of the 10 scrambled images, and tested on the remaining 4 faces and 2 scrambled exemplars.

To assess similarities between responses across stimulus duration conditions, face cross-decoding was also performed, whereby a decoder was trained on 150 ms faces and tested on 30 ms faces and vice-versa. The analysis was repeated for all pairs of conditions, using cross-exemplar cross-validation to ensure true generalization of responses; the resulting accuracies were averaged across the two training/testing directions, which led to similar results.

The temporal structure of face responses was assessed through temporal generalization decoding (King & Dehaene, 2014). Classifier models were trained on each sampled time point between −0.1 and 0.7 s and tested on all time points in order to evaluate the generalizability of neural patterns over time at each stimulus duration. For this analysis, a cross-exemplar hold-out procedure was used to speed up computation (the training and test sets each consisted of 10 face identities/5 scrambled exemplars).

For expression decoding, classification was separately applied to all pairs of emotional expression conditions for each stimulus duration. As low trial numbers were a limitation of the study design, we increased the power of our analysis by also pooling together trials containing faces shown for 30 ms and 150 ms (which were shown to share representations in the cross-decoding analysis). Performance was evaluated using five-fold cross-exemplar cross-validation. Note that splitting the datasets according to perceptual rating led to largely similar results (Supplementary Figure 3).

To achieve equal class sizes in face decoding, face trials were randomly subsampled (after cross-exemplar partitioning) to match the number of scrambled trials. For expression classification, trial numbers did not significantly differ between conditions after artefact rejection (*F* (1.92, 46.18) = 0.15, *P* = 0.85, *η*^2^ = 0.0062).

### MEG sensor-level analyses

MEG data were analyzed using Matlab (Mathworks, Inc.) and the Fieldtrip toolbox (Oostenveld, Fries, Maris, & Schoffelen, 2011). Prior to analysis, trials containing excessive eye or muscle artefacts were excluded based on visual inspection, as were trials exceeding 5 mm in head motion (quantified as the displacement of any head coil between two sampled time points). Using eyetracker information, we also excluded trials containing saccades and fixations away from stimulus or blinks during stimulus presentation. A mean of 8.71% ±9.4% of trials were excluded based on this procedure.

For all analyses, MEG data were downsampled to 300 Hz and baseline corrected using the 500 ms before stimulus onset. A low-pass filter was applied at 100 Hz and a 50 Hz comb filter was used to remove the mains noise and its harmonics.

To improve SNR (Grootswagers, Wardle, & Carlson, 2017), each dataset was divided into 20 equal partitions and pseudo-trials were created by averaging the trials in each partition. This procedure was repeated 10 times with random assignment of trials to pseudo-trials and was performed separately for the training and test sets.

To improve data quality, we performed multivariate noise normalization (MNN; Guggenmos, Sterzer, and Cichy, 2018). The time-resolved error covariance between sensors was calculated based on the covariance matrix (Σ) of the training set (*X)* and used to normalize both the training and test sets, in order to downweight MEG channels with higher noise levels (Equation 1).

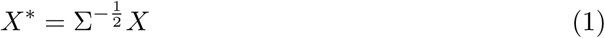

In sensor-level MVPA analyses, all 271 MEG sensors were included as features and decoding was performed for each sampled time point between −0.1 and 0.7 s around stimulus onset.

### MEG source-space analyses

For source analyses, each participant’s MRI (*N* = 24) was coregistered to the MEG data by marking the fiducial coil locations on the MRI and aligning the digitized head shape to the MRI with Fieldtrip. MEG data were projected into source space using a vectorial Linearly Constrained Minimum Variance (LCMV) beamformer (Van Veen, van Drongelen, Yuchtman, & Suzuki, 1997). To reconstruct activity at locations equivalent across participants, a template grid with a 10 mm isotropic resolution was defined using the MNI template brain and was warped to each individual MRI. The covariance matrix was calculated based on the average of all trials across conditions bandpass-filtered between 0.1 and 100 Hz; this was then combined with a single-shell forward model to create an adaptive spatial filter, reconstructing each source as a weighted sum of all MEG sensor signals (Hillebrand, Singh, Holliday, Furlong, & Barnes, 2005). To alleviate the depth bias in MEG source reconstruction, beamformer weights were normalized by their vector norm (Hillebrand, Barnes, Bosboom, Berendse, & Stam, 2012). To improve data quality, MNN was included in the source localization procedure, by multiplying the normalized beam-former filters by the error covariance matrix to ensure that sensors with higher noise levels were downweighted. Next, the sensor-level data were multiplied by the corresponding weighted filters in order to reconstruct the time-courses of virtual sensors at all locations in the brain. This resulted in three time-courses for each source, containing each of the three dipole orientations, which were concatenated for use in the MVPA analysis in order to maximize classification performance (Gohel, Lim, Kim, Kwon, & Kim, 2018). Preprocessing (baseline correction and downsampling) was performed as for sensor-level analyses.

A searchlight approach was used in source-space classification, whereby clusters with a 10 mm radius were entered separately into the decoding analysis. To exclude sources outside the brain and in the cerebellum, we restricted our searchlight analysis to sources included in the 90-region Automated Anatomical Labelling (AAL) atlas (Tzourio-Mazoyer et al., 2002). Given the 10 mm resolution of our sourcemodel, this amounted to a maximum of 27 neighbouring sources being included as features (mean 26.9, median 27, SD 0.31). Source-space subliminal face decoding was performed on 30 ms time windows with a 3 ms overlap using the time windows identified in sensor-space decoding in order to reduce computational cost. We also performed supraliminal face decoding (150 ms faces vs. scrambled stimuli) in order to identify a face-responsive ROI for use in the RSA analysis. This was accomplished by identifying searchlights achieving a cross-subject accuracy above the 99.5th percentile (*P* < 0.005, 66 searchlights; Figure 1).

**Figure 1:**
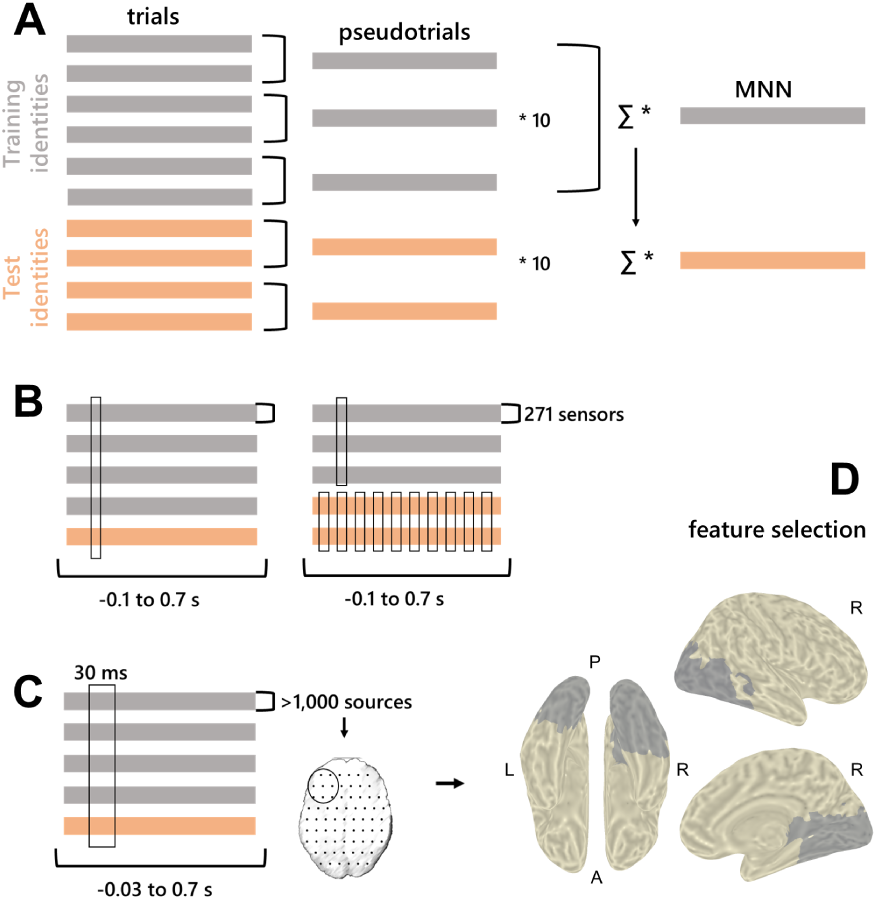
Overview of the MVPA analysis pipeline.**A.** Trial averaging and multivariate noise normalization (MNN) procedure.Σis the error covariance matrix. **B.** Sensor-space time-resolved decoding (left) and temporal generalization (right). **C.** Source-space searchlight decoding procedure. **D.** Sources included in the representational similarity analysis based on face vs. scrambled classification results. P: posterior; A: anterior; L: left; R: right.

### Significance testing

We evaluated decoding performance using the averaged accuracy across subjects (proportion correctly classified trials) and assessed its significance through randomization testing (Nichols & Holmes, 2001).

For sensor-level decoding, 1,000 label shuffling iterations across the training and test sets were used to estimate the null distribution using the time point achieving maximum average accuracy in the MVPA analysis (Dima, Perry, & Singh, 2018). Omnibus correction for multiple comparisons was applied across tests, time points and sources where applicable (Nichols & Holmes, 2001; Singh, Barnes, & Hillebrand, 2003), with a supplementary false discovery rate correction applied for tests where the null distribution was not separately estimated. To avoid spurious effects, a threshold of 5 consecutive significant time points (5^2^ in 2D temporal generalization maps) was imposed. For source-space decoding, 100 randomization iterations were performed for each source cluster and subject in order to reduce computational cost, which were randomly combined into 1000 whole-brain group maps (Stelzer, Chen, & Turner, 2013). A minimal extent of three consecutive time windows with a FDR-corrected *P* < 0.005 was applied.

### Representational Similarity Analysis (RSA)

#### Neural patterns and analysis framework

To interrogate the content of neural representations in space and time, we performed representational similarity analysis (RSA). For this analysis, MEG data were source reconstructed as described above and trials were sorted according to expression and face identity. RSA was performed separately for each stimulus duration and only trials containing faces were included in the analysis. We tracked representational dynamics using a searchlight analysis restricted to the occipitotemporal sources identified in face decoding, with a temporal resolution of 30 ms. The exclusion of responses to scrambled stimuli from the RSA ensured that feature selection was based on an orthogonal contrast (Figure 1).

To create MEG representational dissimilarity matrices (RDMs), we calculated the squared cross-validated Euclidean distance between all pairs of face stimuli (Guggenmos et al., 2018). Note that as the data were multivariately noise-normalized, this is equivalent to the squared cross-validated Mahalanobis distance (Walther et al., 2016). For each participant, the data were split into a training set (the first 2 sessions) and a test set (the last 2 sessions). The two stimulus repetitions contained in each set were averaged, and these were averaged across subjects to create training and test sets. To compute the cross-validated Euclidean distance between two stimulus patterns (*X*^∗^, *Y*^∗^), we calculated the dot products of pattern differences based on the training set and the test set (Equation 2). This procedure has the advantage of increasing the reliability of distance estimates in the presence of noise.

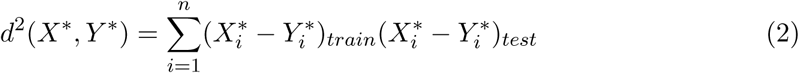

The spatiotemporally resolved MEG RDMs were then correlated with several model RDMs to assess the contribution of different features to neural representations. In an initial analysis, we calculated Spearman’s rank correlation coefficients between each model RDM and the MEG RDM (Nili et al., 2014). To further investigate the unique contribution of each model, we entered the significantly correlated models based on visual features of the images into a partial correlation analysis, where each model’s correlation to the MEG data was recalculated after partialling out the contribution of the other models.

Note that a model based on behaviour, which was also represented in the MEG data for all stimulus duration conditions, was not included in the partial correlation analysis; the rationale is that we were interested in the contribution of each visual property independently of the others, but we did not expect a unique contribution of behaviour in the absence of expression-related visual properties, and partialling out the behavioural model from the visual models would not be easily interpretable. Instead, we preferred to independently describe the correlations between behaviour and visual features, brain and behaviour, and brain and visual features, as the three main factors of interest in our analysis.

#### Model RDMs

We investigated the temporal dynamics of face perception by assessing the similarity between MEG patterns and 9 models quantifying behaviour and facial/visual properties (Figure 2).

**Figure 2:**
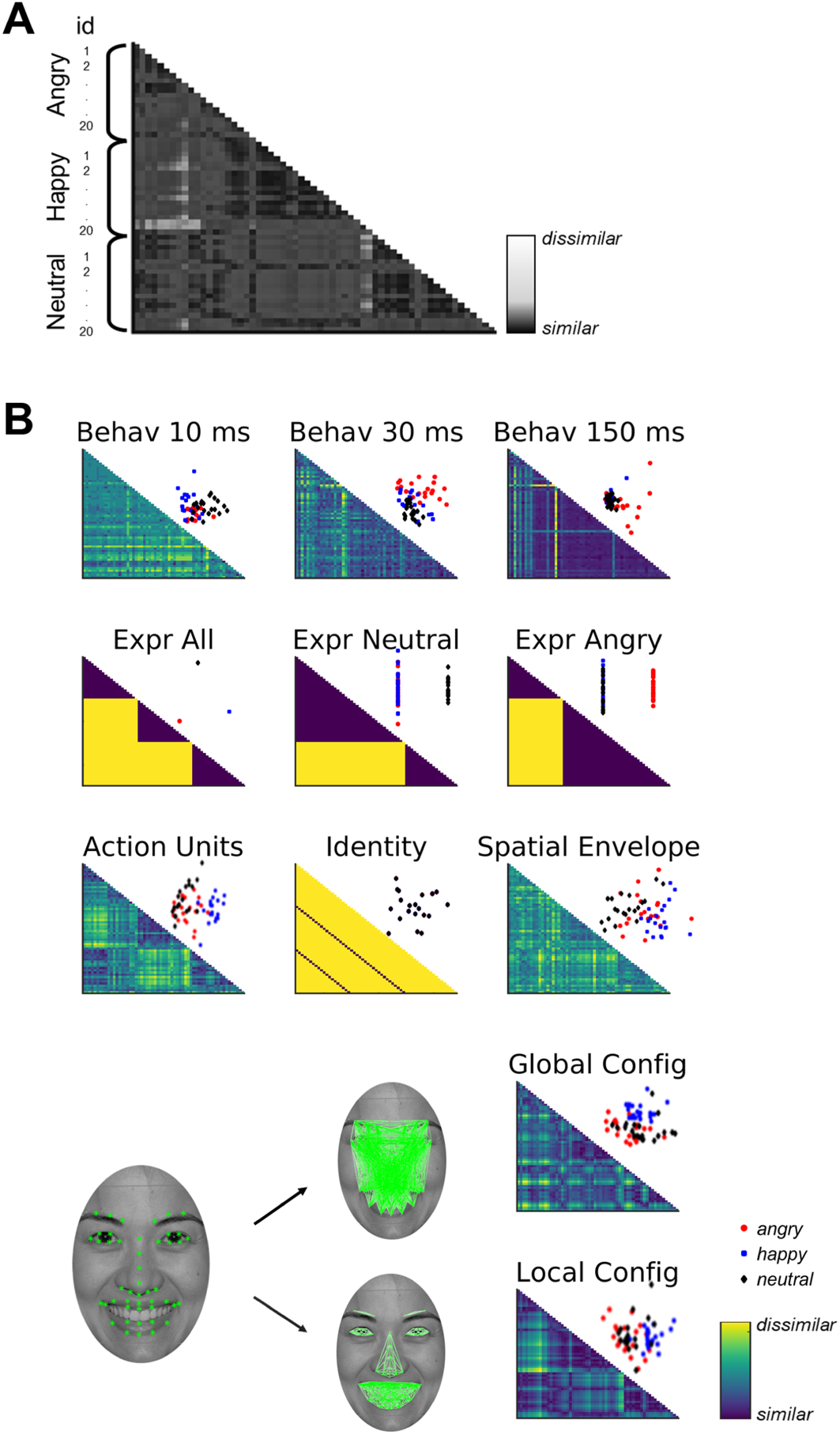
Models used in RSA analysis. **A.** Example model RDM: each model maps pair-wise dissimilarities between faces, which are sorted according to expression and identity. **B.** Model RDMs showing predicted distances between all pairs of stimuli. 2D multidimensional scaling (MDS) plots are shown above each model to visualize how the three expression categories are organized according to each model. For the local and global configuration models, we also show the facial landmarks and the within-feature/between-feature distances used to create each model. Behav: behavioural models; Expr: high-level expression models (all-vs-all, neutral-vs-others, and angry-vs-others); Config: face configuration models.

To create behavioural model RDMs, we calculated the number of error responses made by each participant to each stimulus and summed these up to create a cross-subject behavioural RDM. For each stimulus duration, we created separate behavioural RDMs by calculating pairwise cross-validated Euclidean distances between error response patterns, using a cross-session training/test split as described above.

To create face configuration RDMs, we first used OpenFace (Baltrusaitis, Robinson, & Morency, 2016) to automatically detect and label face landmarks. The software created 68 2D landmarks for each face. We removed landmarks corresponding to the face outline and the 2 outermost eyebrow landmarks, to account for cases in which these landmarks were cropped out by the oval mask used in the MEG stimulus set. The final landmark set consisted of 47 coordinates for 6 facial features (eyes, eyebrows, nose, and mouth), which were visually inspected to ensure that they were correctly marked. To capture feature-based (local) facial configuration, we calculated within-feature pairwise Euclidean distances between landmarks (Figure 2B). To quantify global face configuration, we calculated between-feature Euclidean distances (the distances between each landmark and all landmarks belonging to different facial features). Distances were then concatenated to create feature vectors describing each face in terms of its local/global configuration, and Euclidean distances between them gave the final configural model RDMs. These models correspond to the featural and configural stages in classic models of face perception (Diamond & Carey, 1986; Piepers & Robbins, 2012).

To create a high-level identity model, we assigned distances of 0 to pairs of face identities repeated across emotional expression conditions, and distances of 1 to pairs of different face identities. We used a similar strategy to create high-level emotional expression models. An all-versus-all model was created by assigning distances of 0 to all faces belonging to the same emotional expression condition, and distances of 1 to pairs of faces differing in emotion. We also tested a neutral-versus-others model by assigning distances of 0 to all emotional faces (happy + angry), and an angry-versus-others model by assigning distances of 0 to all benign faces (happy + neutral).

To account for variability in expression that is not captured by such high-level binary representations, we also tested a model based on Action Units. Action Units quantify changes in expression by categorizing facial movements (Ekman & Friesen, 1977). We used OpenFace (Baltrusaitis et al., 2016) to automatically extract the intensity of 12 Action Units in our image set (Supplementary Table 4), and we calculated pairwise Euclidean distances between these intensities for all pairs of faces in our stimulus set to obtain an Action Unit RDM.

Finally, a spatial envelope model was created in order to capture image characteristics using the GIST descriptor (Oliva & Torralba, 2001). This procedure extracts 512 values per image by applying a series of Gabor filters at different orientations and positions, and thus quantifies the average orientation energy at each spatial frequency. To obtain the spatial envelope RDM, we calculated pairwise Euclidean distances between all images using the GIST values.

Finally, models were subject to multidimensional scaling (MDS) to visualize how each model represents the similarity between facial expressions in a 2D space (Figure 2).

#### Significance testing

To assess the significance of spatiotemporally resolved correlation maps, we used a randomization approach. Model RDMs were shuffled 1,000 times and correlations were re-computed for each of the 66 searchlights using the time window achieving the maximal correlation coefficient across models for each of the stimulus duration conditions. Since negative correlations were not expected and would not be easily interpretable, P-values were calculated using a one-sided test (Furl, Lohse, & Pizzorni-Ferrarese, 2017). To correct for multiple comparisons, P-values were omnibus-corrected by creating a maximal distribution of randomized correlation coefficients across searchlights, models and conditions, and FDR and cluster-corrected across timepoints (*α* = 0.05, thresholded at 3 consecutive time windows).

#### Variance partitioning

To gain more insight into the relationship between behavioural responses, expression categories and face configuration models, we used a variance partitioning approach (Greene, Baldassano, Esteva, Beck, & Fei-fei, 2016; Groen et al., 2018). For each stimulus duration condition, the corresponding behavioural RDM was entered into a hierarchical multiple linear regression analysis, with three model RDMs as predictors: the two facial configuration models and the most correlated high-level expression model (10 ms: neutral-vs-others; 30 and 150 ms: angry-vs-others). These models were selected to reduce the predictor space before performing variance partitioning. To quantify the unique and shared variance contributed by each model, we calculated the *R*^2^ value for every combination of predictors (i.e. all three models together, each pair of models separately, and each model separately). The EulerAPE software was used for visualization (Micallef and Rodgers, 2014; Figure 6B).

## Results

### Perception and behaviour

We assessed the effects of stimulus duration and face expression on behaviour using a 3 × 3 repeated-measures ANOVA with factors *Duration* (levels: 10 ms, 30 ms, 150 ms) and *Expression* (levels: angry, happy, neutral) on rationalized arcsine-transformed accuracies (see Methods). Stimulus duration had a strong effect on expression discrimination performance, with average performance not exceeding chance level at 10 ms (33.45% ± 2.99) and rising well above chance at 30 and 150 ms (78.62% ± 2.11 and 91.83% ± 1 respectively). This was reflected in a significant main effect of duration in the ANOVA (*P* < 0.0001, *F* (1.21, 29.06) = 221.05, *η*^2^ = 0.9). Face expression had a weak effect, with angry faces categorized less accurately than both happy and neutral faces (*P* = 0.046, *F* (1.95, 46.71) = 3.33, *η*^2^ = 0.12), and with no significant interaction effect (*P* = 0.23, *F* (1.74, 41.83) = 1.53, *η*^2^ = 0.06).

Participants found the task challenging, as reflected in the perceptual awareness ratings: 84.5% of the 10 ms trials were rated as not containing a face (Figure 3E). This suggests that participants were complying with the task with respect to both expression discrimination and perceptual rating. Importantly, for faces presented for 10 ms, there was no difference in accuracy between expressions (*P* = 0.43, *F* (1.65, 39.5) = 0.8) or between any pair of cells in the confusion matrix (*P* = 0.6, *F* (3.42, 82.07) = 0.64), suggesting that faces presented at this duration were equally likely to be categorized as any expression.

**Figure 3:**
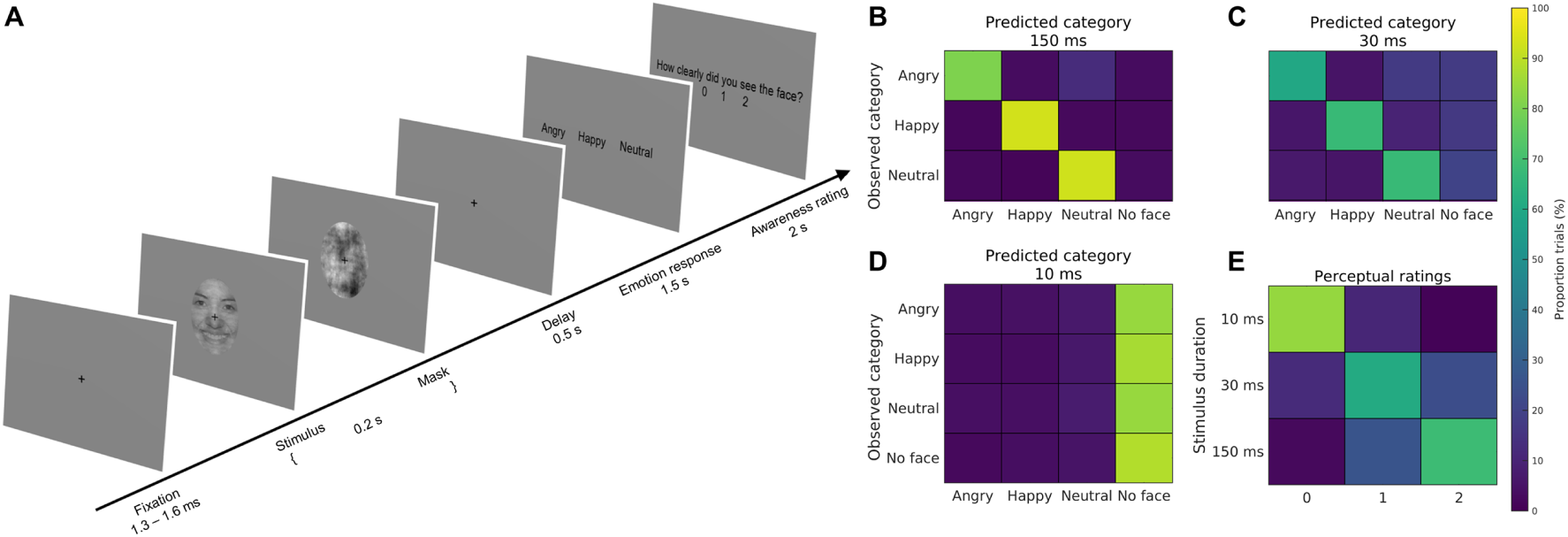
Overview of the experimental paradigm and behavioural results. **A.** Stimuli were presented on screen for 150 ms, 30 ms, or 10 ms, and were followed by a 50 ms, 170 ms, or 190 ms scrambled mask. **B-D.** Confusion matrices mapping the average proportion of trials receiving each of the possible responses (X-axis) out of the trials belonging to each category (Y-axis). “No response” trials were excluded for statistical analysis, but are shown here as representing a “no face” response. **E.** Perceptual ratings for each stimulus duration summarized as average proportion of trials.

### Spatiotemporal dynamics of face perception

To investigate face processing as a function of stimulus duration, we performed within-subject cross-identity decoding of responses to faces vs. scrambled stimuli. The analysis included three components: sensor-level time-resolved classification to evaluate the progression of condition-related information; sensor-level temporal generalization to assess the temporal structure of this information; and source-space decoding to obtain spatial information about subliminal responses to faces (Figure 1).

We first decoded responses to neutral faces vs. scrambled stimuli using data from all MEG sensors, separately for each stimulus duration. In the case of faces presented for 10 ms, any trials reported as containing a face were excluded, to ensure that we assessed responses outside of subjective awareness. Scrambled stimuli could be discriminated from faces presented for 150 and 30 ms starting as early as 100 ms (Figure 4A). After the initial peak in performance, decoding accuracy decreased, but remained well above chance for the remainder of the decoding time window. For faces presented for 10 ms and reported as not perceived, there was only a weak increase in decoding performance, which reached significance at 147 ms and dropped back to chance level after ∼350 ms (Supplementary Table 1). To assess how well face representations generalized across stimulus durations, we repeated this analysis by training and testing on stimulus exemplars presented for different amounts of time (Figure 4B). Decoding accuracy was high when cross-decoding between 30 ms and 150 ms faces, with two increases in performance at M170 latencies (100-200 ms) and after 300 ms. On the other hand, representations only generalized to 10 ms faces for a limited time window corresponding to the M170 component.

**Figure 4:**
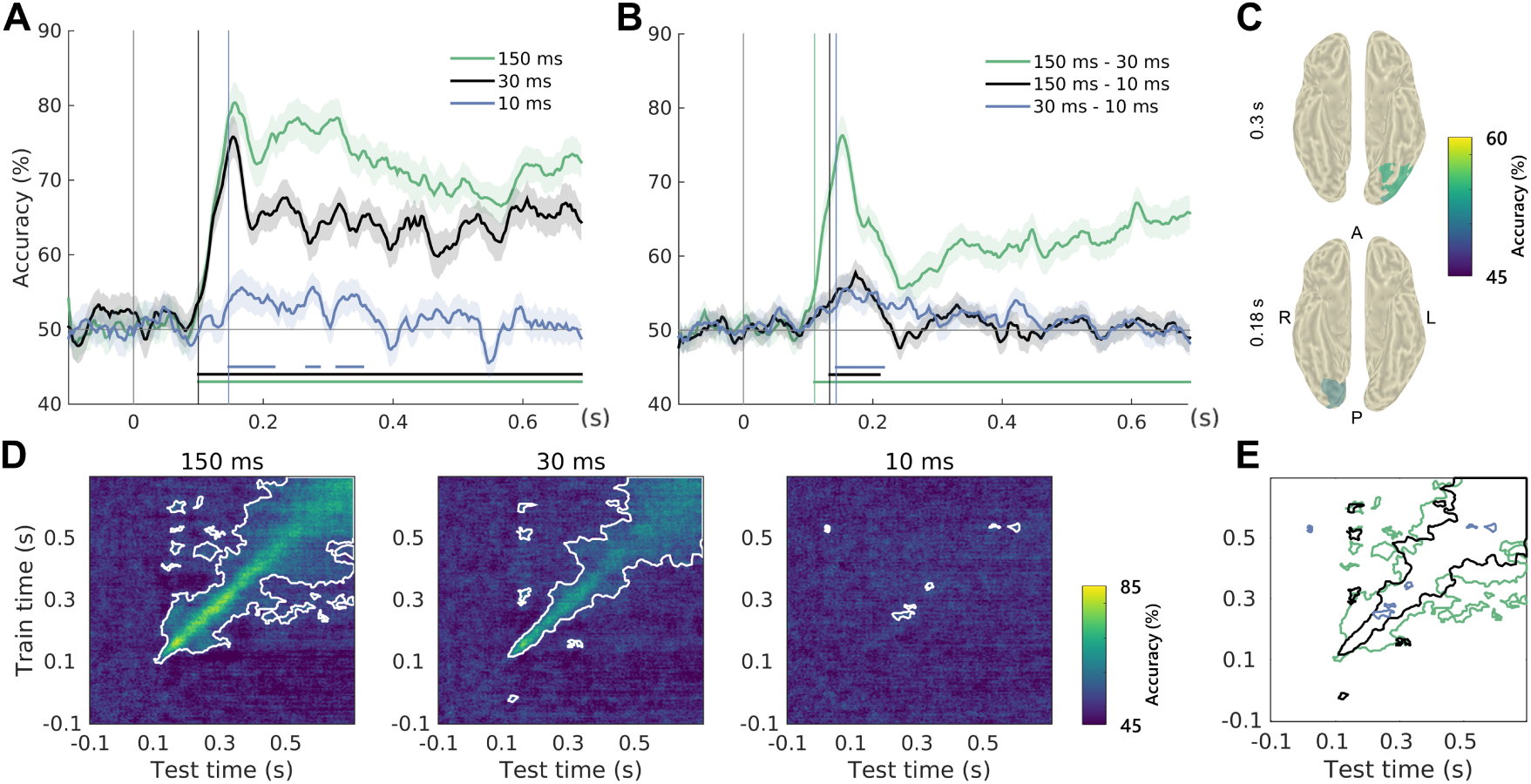
Face vs. scrambled decoding results. **A.** Sensor-space time-resolved decoding accuracy for all stimulus durations. Colour-coded vertical bars mark above-chance decoding onset and horizontal lines show significant time windows (*P* < 0.05, corrected). **B.** Sensor-space time-resolved cross-decoding for all pairs of stimulus durations. Cross-validation was performed across exemplars and accuracies were averaged over the two training/test directions. **C.** Sources achieving above-chance decoding of 10 ms faces out-side awareness at M170 latencies in source space (*P* < 0.005, corrected). **D.** Sensor-space temporal generalization accuracy and significant clusters (white contours; *P* < 0.05, corrected) for all stimulus durations. **E.** Significant temporal generalization clusters for all three stimulus durations, showing more sustained representations of faces presented for 150 ms (legend as in A).

Using temporal generalization decoding (King & Dehaene, 2014), we investigated the temporal structure of face responses, and we found that this changed with stimulus duration. For faces presented for 150 ms, successful temporal generalization started at ∼93 ms in a diagonal pattern suggestive of transient representations, with more sustained representations (square patterns) arising at M170 latencies and after 300 ms (Figure 4D-E). For 30 ms stimuli, a transient representation pattern started at ∼110 ms after stimulus onset and sustained representations only arose later (∼400 ms). Early processing thus appears to be heavily biased by stimulus presentation duration, with 30 ms faces failing to elicit a stable representation at M170 latencies. For faces presented for 10 ms, only few transient clusters survived correction for multiple comparisons, with the largest one occurring after 200 ms.

Finally, we spatially localized the subliminal response to faces in source space. All participants with one exception acquired a structural MRI, which was used to source localize the MEG data using a Linearly Constrained Minimum Variance (LCMV) beamformer (Van Veen et al., 1997). We performed whole-brain searchlight classification of 10 ms faces vs. scrambled stimuli (*N* = 24), using source clusters with a radius of 10 mm and time windows of 30 ms. Faces were successfully decoded in a right occipital area at M170 latencies (Figure 4C), with a later stage associated with ventral patterns.

### Temporal dynamics of expression perception

Next, we performed sensor-level cross-identity decoding of all pairs of emotional expressions separately for each stimulus duration. The analysis was performed similarly to the time-resolved face decoding analysis described above.

The highest decoding performance was achieved on late responses to expressions presented for 150 ms (Figure 5A). Expressions presented for 30 ms also achieved above-chance decoding, although these effects were more transient. We also performed this analysis on pooled datasets (faces presented for 30 and 150 ms), as the face cross-decoding analysis showed that responses generalized between these two categories (Figure 4B). This revealed a multi-stage progression for all expressions, with transient early decoding at M100 latencies and an increasing accuracy at later stages (Figure 5B). We found no above-chance performance when decoding 10 ms expressions. This finding adds to emerging evidence against the automatic processing of expression outside awareness (Koster, Verschuere, Burssens, Custers, & Crombez, 2007; Pessoa, Japee, & Sturman, 2006; Hedger, Gray, Garner, & Adams, 2016; Schlossmacher, Junghöfer, Straube, & Bruchmann, 2017), and we explore potential reasons for this result below.

**Figure 5:**
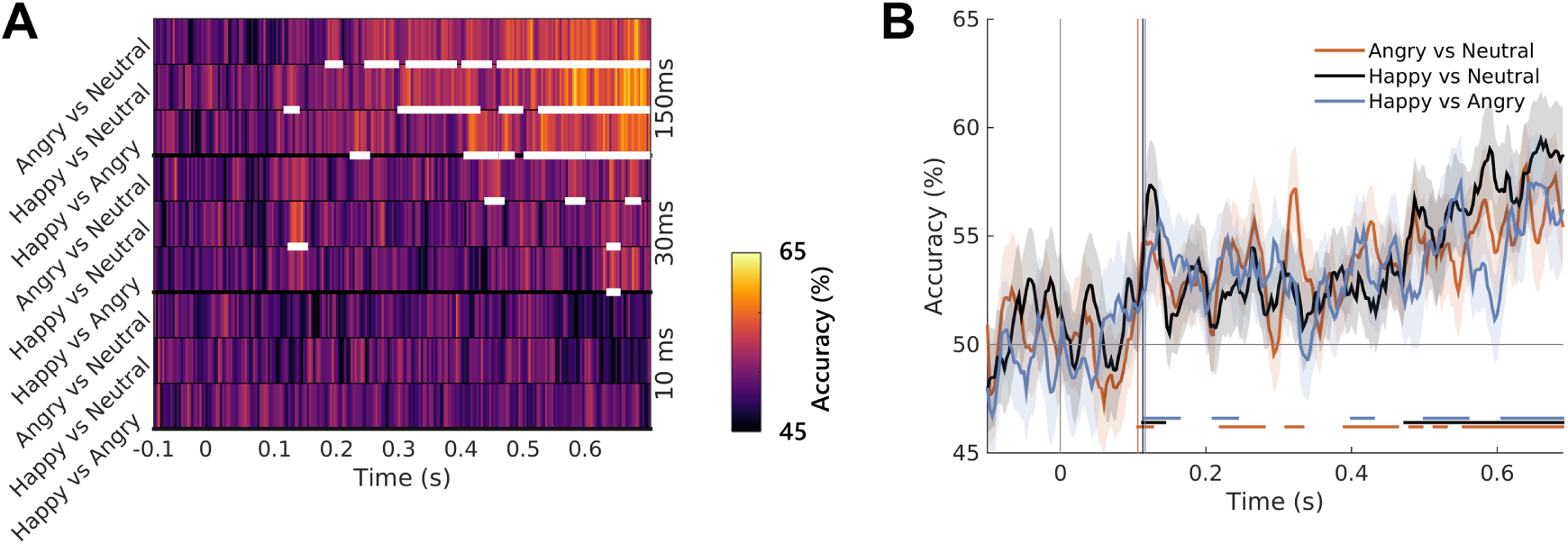
Expression decoding results. **A.** Time-resolved decoding accuracy for the three expression decoding problems and the three stimulus durations. White horizontal lines show significant time windows (*P* < 0.05, corrected). **B.** Time-resolved accuracy for the three expression decoding problems using the pooled datasets (30 + 150 ms).

### Face representations in occipitotemporal cortex

To interrogate the neural representations underpinning these pattern differences, we performed representational similarity analysis (RSA) using a searchlight approach at the source level (Su, Fonteneau, Marslen-wilson, & Kriegeskorte, 2012) in a face-responsive area of interest determined using an orthogonal contrast. We investigated the representational dynamics of face perception by assessing the similarity between MEG patterns and models quantifying behaviour, face features, face configuration, expression, identity and visual properties, using both a Spearman’s rank correlation (Nili et al., 2014) and partial correlation (see Methods).

### Occipitotemporal cortex encodes behavioural responses

Among the other model RDMs tested, behavioural RDMs correlated most with the high-level expression models (particularly the angry-vs-others model at 30 ms and 150 ms, Spearman’s *ρ* = 0.29 and *ρ* = 0.34). At 150 ms, the behavioural RDM also correlated with the configural face models (*ρ* = 0.22 and *ρ* = 0.18). As expected based on performance, behavioural RDMs at 10 ms did not correlate with the other two (*ρ* = −0.05 and *ρ* = −0.09 respectively), while behavioural RDMs at 30 and 150 ms were positively correlated (*ρ* = 0.38; Figure 6A).

**Figure 6:**
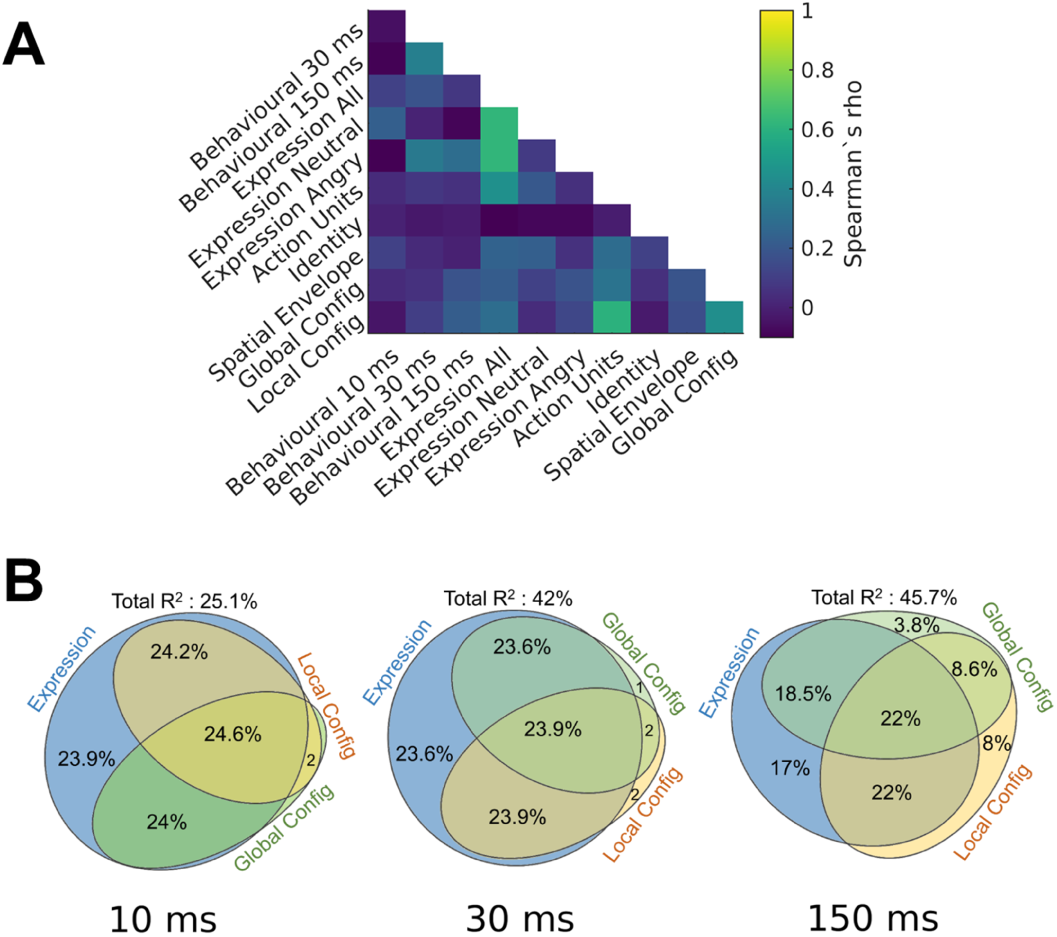
Relating behaviour to representational models. **A.** Model inter-correlations (Spearman’s *ρ*). **B.** Variance partitioning results, showing the contributions of expression and face configuration models to behavioural responses at each stimulus duration. Values represent % of the total *R*^2^.

Based on these links, face configuration, together with facial expression, appears to partially explain behavioural responses. To test this, we performed a variance partitioning analysis, using hierarchical multiple regression to quantify the unique and shared variance in behaviour explained by facial configuration and high-level expression models. In the 10 ms condition, the neutral-vs-others model and the two configural models explained 25.1% of the variance; in the 30 ms and 150 ms conditions, the angry-vs-others model and the configural models explained up to 45.7% of the variance in behaviour. Furthermore, while the expression model contributed most of the variance, over 75% of this variance was shared with the configural models. The unique contribution of the configural models increased with stimulus duration (from ∼2% at 10 ms, to ∼20% at 150 ms). Together, these results point to the role of face configuration in driving high-level representations and behaviour. Note that we were unable to decode 10 ms expressions from the MEG data; however, the variance partitioning analysis of behavioural responses in this condition showed a contribution of both facial expression and configuration to behaviour.

Behavioural RDMs showed the strongest and most sustained correlations with MEG patterns in ventral stream areas, including sources corresponding to the location of the fusiform face area (FFA) and occipital face area (OFA; Figure 7). Behavioural representations evolved differently in time for the three stimulus durations. For 10 ms faces, behaviour explained the data starting at 120 ms until the end of the analysis time window. Representations emerged similarly early for 150 ms faces and reached the noise ceiling before decreasing again at 400 ms. For 30 ms faces, correlations were significant starting at 210 ms in a relatively focal right temporal area. Patterns were more posterior for 10 ms faces and more extensive, including sources corresponding to the OFA and FFA, for 150 ms faces.

**Figure 7:**
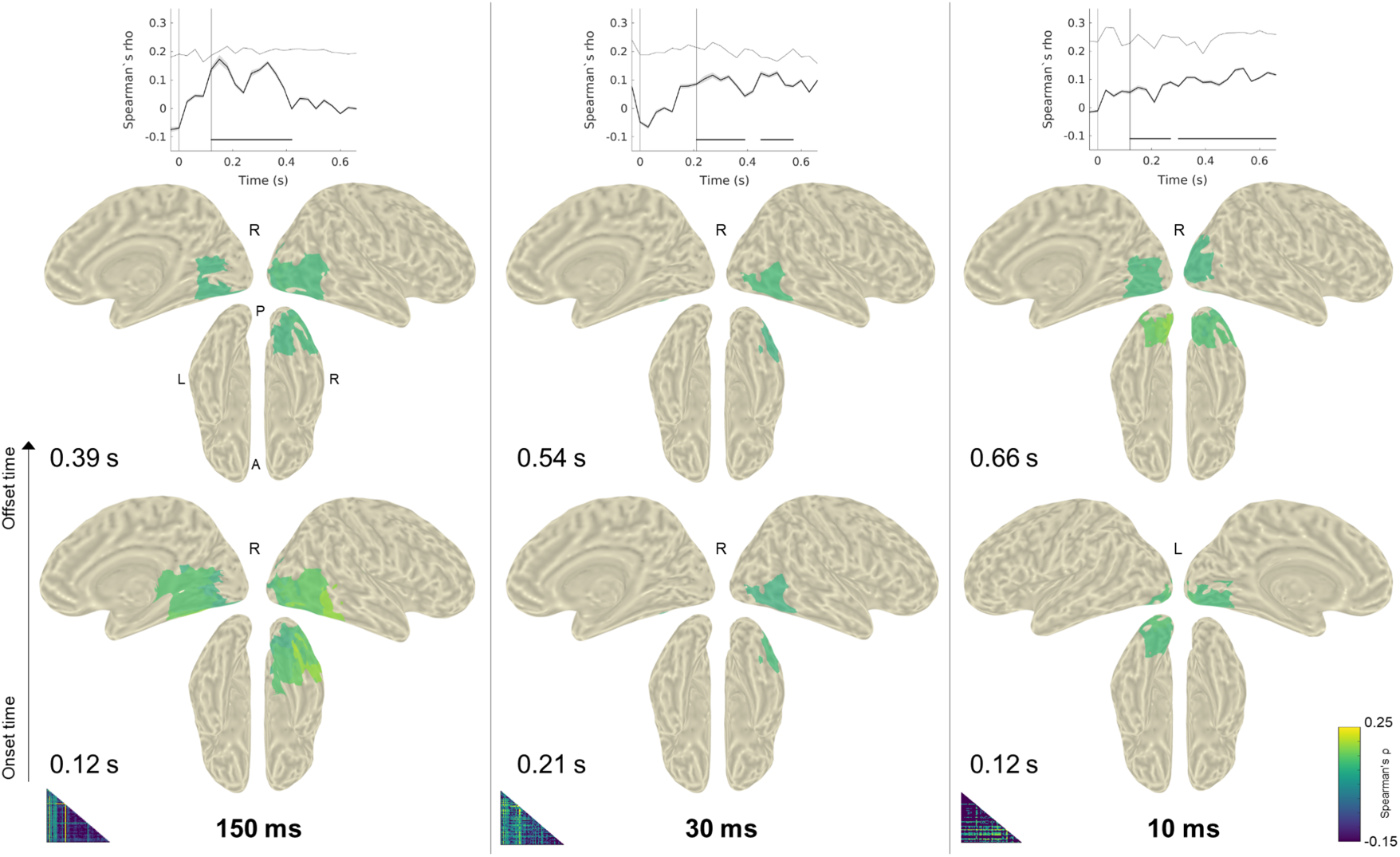
Correlations between MEG patterns and behavioural model RDMs for each stimulus condition duration (vertical columns). The top panels show correlation time-courses averaged across all significant searchlights; the noise ceiling is shown as a dotted horizontal line and is only approached in the 150 ms condition. The cortical maps show significant correlation coefficients for the first and last significant time windows (onset and offset times) on the inflated template MNI brain. The hemisphere shown is indicated with the letter R/L. Model RDMs are shown in the lower left corner of each column. See SourceMovies1 for movies showing the evolution of behavioural representations in time.

The correlation time-courses suggest interesting differences in processing as a function of the information available: for clearly perceived faces, features relevant in behaviour are extracted between 120-400 ms, while behavioural responses for briefly presented faces appear to require sustained processing, as reflected by behaviour-related correlations not dropping back to zero. These results are in line with previous evidence of behavioural representations in ventral stream areas in scene and object perception (Walther, Caddigan, Fei-Fei, & Beck, 2009), and suggest that visual feature processing, even at early stages, is closely linked to behavioural goals.

### Configural face processing from featural to relational

The two face configuration models were also represented in the MEG patterns. In the correlation analysis, the local and global configuration models explained representations in partially overlapping areas of the ventral stream (corresponding to the right FFA location), with local configuration representations arising earlier (at 120 ms for 150 ms faces, and 360 ms for 30 ms faces). The RSA method used here favoured sustained correlations over transient peaks; note that the global configuration model approached the noise ceiling during a transient time window at M170 latencies for both 150 ms and 30 ms faces, suggesting a contribution of second-order characteristics, although this occurred later than feature representations (Supplementary Figure 4). The partial correlation analysis revealed further differences between conditions: for 150 ms faces, the local and global models made unique, successive contributions in explaining the data; conversely, for 30 ms faces we detected no unique contributions, suggesting that the extraction of configural information from faces occurs differently in the absence of sufficient information. None of the models significantly correlated with MEG patterns elicited by 10 ms faces.

Note that although both internal (eyes, nose, mouth) and external (face shape, hair) face features have been shown to contribute to neural responses to faces (Axelrod, 2010), we focus here on internal features; for the purposes of this paper, external features were excluded from the stimuli and we refer to the second-order configuration of distances between internal features as “global configuration”. Internal features are relevant to the context of expression discrimination and have been shown to be more reliable even in facial recognition contexts (e.g. Longmore, Liu, and Young, 2015).

### Transient representations of visual and high-level models

Two other models elicited brief representations in the MEG data. For 150 ms faces, the spatial envelope model explained left hemisphere occipital representations starting at ∼400 ms, suggesting sustained processing of visual features, potentially based on feedback mechanisms. For 30 ms faces, a high-level expression model (neutral-vs.-others) was represented in the MEG data starting at 300 ms (Figure 9). This can be speculatively explained by the formation of task-related representations in the absence of sufficient information. On the contrary, when faces are clearly presented, only models encoding face characteristics are represented, while categorical models show no contribution to occipitotemporal representations. Note that despite the role of facial features in explaining neural responses, the Action Unit model RDM did not significantly correlate with the MEG patterns, probably due to the static and brief nature of our stimuli.

Although correlation coefficients between the models and neural data are generally low (Supplementary Table 3), the noise ceiling shows that the maximal correlation possible with our data is also low (mean *ρ*=0.21); this is not surprising, considering the low *ρ*-values usually found in MEG RSA studies, and the fact that our paradigm involved complex, high-level visual stimuli and a demanding task. In this case, the noise ceiling serves as a useful benchmark for the explanatory power of our models. For example, the behavioural RDM reaches the noise ceiling in the 150 ms condition, but not for briefer stimuli, suggesting that behavioural representations fully explain the data when stimuli are clearly perceived. The local configuration model also shows good explanatory power at its earliest stage, and the same is true for the global model for a brief time window. Other significant models do not reach the noise ceiling (Supplementary Figure 4); given the complex face processing and task-related activity reflected by the MEG patterns, this is not surprising. In fact, the explanatory power of the configural models at early stages (100-200 ms) is striking, as is the strength of behavioural representations in ventral stream within 400 ms. Furthermore, the initial peak in performance of the behavioural model overlaps with the peak of the local configuration model. Together with the shared variance between configuration, expression and behaviour shown in the variance partitioning analysis (Figure 6D), this points to the role played by facial configuration in the extraction of emotional cues essential in the expression discrimination task.

## Discussion

The cross-identity decoding and representational similarity analyses described here converge to highlight the dynamic nature of face representations in the ventral visual stream. Face feature and face configuration representations link occipitotemporal neural patterns and behavioural responses during an expression discrimination task, while their temporal dynamics change to accommodate challenging viewing conditions.

In the time-resolved decoding analysis, a response to faces (150 ms and 30 ms) emerged at ∼100 ms, while faces shown outside of subjective awareness were decodable for a brief time window (147 −350 ms), in line with previous studies showing evidence of face perception outside of awareness (Axelrod, Bar, & Rees, 2015). Temporal representations also varied with stimulus duration: for 150 ms faces, a sustained representation emerged at M170 latencies which was absent for 30 ms faces. This suggests that clearly presented faces are perceived through a multi-stage process, while disrupted recurrent processing leads to delayed stable representations. Although the M170 component decreases in amplitude with face duration (Supplementary Figure 1), its duration does not predict such a marked change in temporal structure, especially given the high decoding accuracy at this latency obtained in both conditions in the time-resolved face decoding analysis. Trial-to-trial variability, cited as another potential explanation for diagonal patterns (Vidaurre, Myers, Stokes, Nobre, & Woolrich, 2018), is also not expected to systematically vary between our conditions. On the other hand, sustained representations in temporal generalization analyses are thought to be reflective of conscious perception and recurrent processes (Dehaene, 2016). It has previously been suggested that faster stimulus presentation leads to more transient representations (Mohsenzadeh, Qin, Cichy, & Pantazis, 2018); however, the backward masking procedure used here disrupts the formation of a stable representation by entering the visual stream, and it is unclear whether different methods of preventing awareness would lead to the same results.

Information supporting face decoding outside of subjective awareness was localized to occipitotemporal cortex in our searchlight source-space decoding analysis (Figure 4C). Given the disruption of recurrent processing in backward masking (Lamme, Zipser, & Spekreijse, 2002; Boehler, Schoenfeld, Heinze, & Hopf, 2008), the early stages of this response can be attributed to either purely feedforward activity, or to feedback connections targeting V1 at early processing stages (Wyatte, Jilk, & O’Reilly, 2014; Mohsenzadeh et al., 2018). Furthermore, the fact that we detect a response to faces, and not to expression, suggests that the different tasks of identification and categorization are supported by qualitatively different mechanisms. However, the spatial resolution of MEG, together with recent observations of information spreading in searchlight source-space MVPA (Sato, Yamashita, Sato, & Miyawaki, 2018), prevent us from drawing strong conclusions about the origin of this response to faces. To minimize such concerns, we restricted our source-space decoding analysis to localizing effects identified at the sensor level, and we applied randomization testing with an omnibus threshold in order to avoid spurious effects.

All expressions presented for at least 30 ms were decodable from MEG data starting at ∼100 ms. Since all analyses were performed across facial identity and stimuli were matched for low-level properties, this suggests that expression categorization begins at the early stages of visual perception (Aguado et al., 2012; Dima, Perry, Messaritaki, Zhang, & Singh, 2018), in line with behavioural goals. However, in terms of non-conscious expression processing, the results are mixed. Despite the absence of a subliminal expression effect in MEG responses, behavioural data suggest that expression (specifically, a model differentiating between emotional and neutral stimuli) explains approximately one quarter of the variance in behavioural responses given to faces presented for 10 ms. This effect is not revealed by the more traditional accuracy-based behavioural analysis, suggesting that model-based approaches to the analysis of behavioural responses can provide additional information. With the caveat that low numbers of trials were included in this analysis, the fact that cross-subject patterns of response reflected shared variance between the models based on expression, facial features and facial configuration points to a certain degree of expression processing taking place outside of subjective awareness. The absence of a subliminal expression effect in the neural data may be explained by several factors, including the limited ROI used in RSA, the study design minimizing residual awareness, and challenges in the detection of a potential subcortical response.

Representational similarity analysis results linked stages in time-resolved decoding to stages in feature extraction and to behavioural responses. Ventral stream areas encoded sustained and extensive behavioural representations as early as 120 ms after stimulus onset (Figure 7), suggesting that the extraction of features essential in behavioural decision-making is a rapid process accomplished in face-responsive cortex. This is in line with evidence found in higher-level object and scene perception (Walther et al., 2009; Bankson, Hebart, Groen, & Baker, 2018; Groen et al., 2018) and with previous studies showing that the perceptual similarity of faces is represented in neural patterns (Said, Moore, Engell, & Haxby, 2018; Furl et al., 2017).

Furthermore, ventral stream areas encoded facial features prior to facial configuration when faces were presented for 150 ms. This adds to evidence suggesting that emotional face perception is supported by the processing of diagnostic features, such as the eyes and mouth (Wegrzyn, Vogt, Kireclioglu, Schneider, & Kissler, 2017). What is more, configural representations explain behaviour and overlap with behavioural representations, suggesting that it is face configuration that drives expression-selective responses in ventral stream areas and guides behaviour.

Previous studies have shown differential modulation of ERP components by first-order and second-order face configuration. Some studies have shown early components (P1, N170) to encode the former only (Mercure, Dick, & Johnson, 2008; Zion-Golumbic & Bentin, 2007), while others have also shown effects of second-order configuration at N170 latencies (Eimer, Gosling, Nicholas, & Kiss, 2011). Furthermore, fMRI studies have reported a division of labour in the face-selective network, with the FFA thought to play a special role in representing both types of configural information (Golarai, Ghahremani, Eberhardt, & Gabrieli, 2015). Recently, it has been suggested that featural and configural processing of even non-face objects elicit face-like responses in the OFA and FFA (Zachariou, Safiullah, & Ungerleider, 2018). Here, we combined the strengths of source-localized MEG data and the RSA framework to tease apart the two models using a single stimulus set. The searchlight RSA analysis revealed that the two models overlap spatially in a right ventral stream area corresponding to the FFA, but are dissociated temporally: for 150 ms faces, representations switch from first-order to second-order at ∼300 ms after stimulus onset, bringing together previous fMRI and electrophysiological findings.

Furthermore, this two-stage process appears to depend on the amount of information available to the visual system. For 150 ms faces, local and global configuration models make unique, temporally distinct contributions to explaining the data, as shown in the partial correlation analysis. For 30 ms faces, no unique variance is explained by the two models; furthermore, representations are temporally overlapping in the correlation analysis and occur after 300 ms (Figure 8). This complements our sensor-level temporal generalization findings: 30 ms faces are processed through a series of transient coding steps at early stages and a stable representation is formed after 300 ms, when both first-order and second-order features are represented. On the other hand, for 150 ms faces, a two-stage process takes place, with an initial stable representation emerging at M170 latencies and supported mainly by first-order features, and a later representation after 300 ms encoding second-order configuration. Feature representations thus appear to be linked to the late emergence of stable representations, thought to be reflective of recurrent processing and categorization. Importantly, this idea is supported by spatially and temporally overlapping behavioural representations in ventral stream areas.

**Figure 8:**
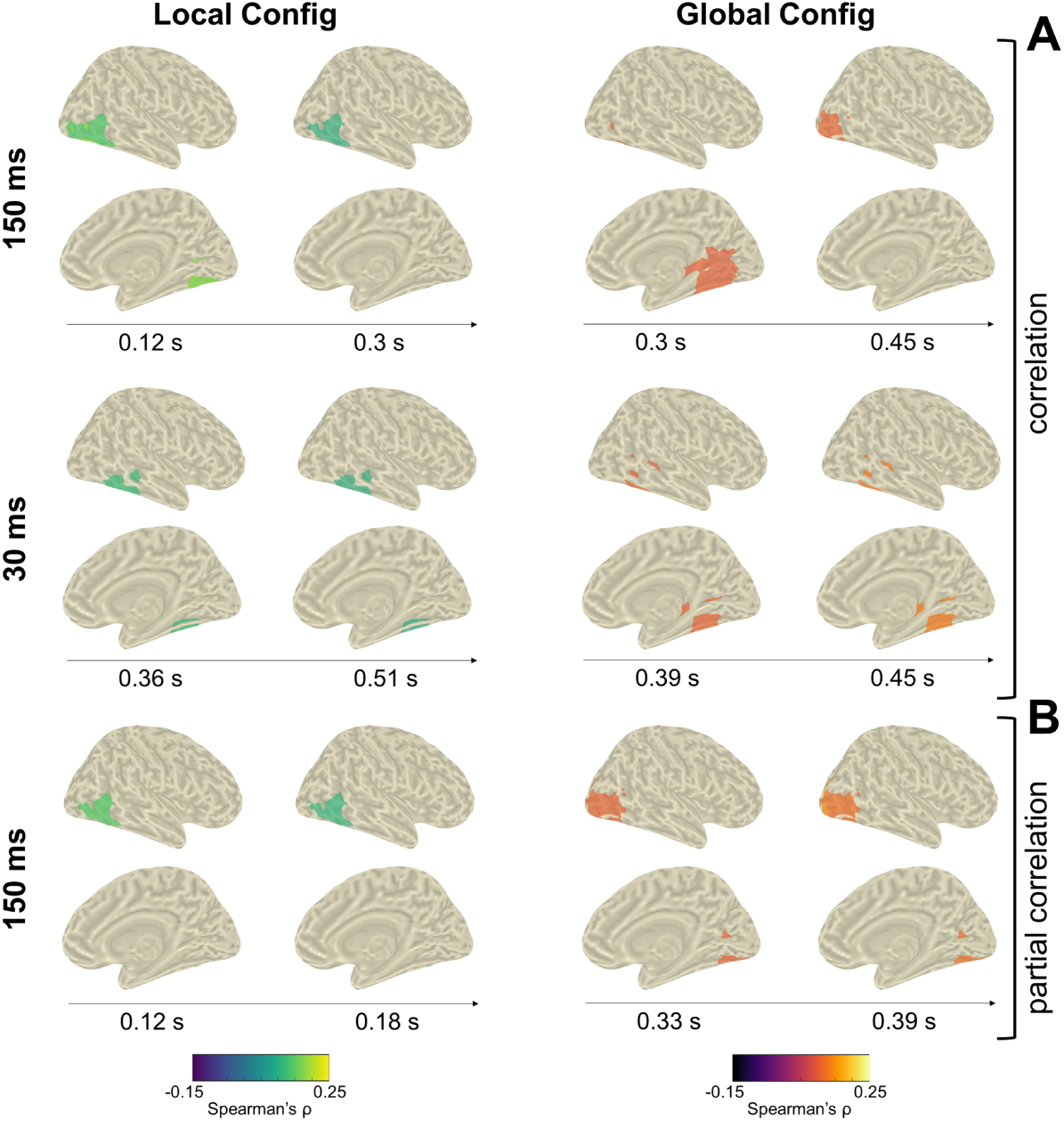
Significant correlations between MEG patterns and configural model RDMs. **A:** Correlation analysis results are significant for the 150 ms and 30 ms conditions. **B:** Partial correlation results are significant for the 150 ms condition. Only right hemisphere searchlights correlate with the configural models. Maps are shown for the onset and offset times of significant correlation. See SourceMovies2 for movies showing the evolution of behavioural representations in time.

**Figure 9:**
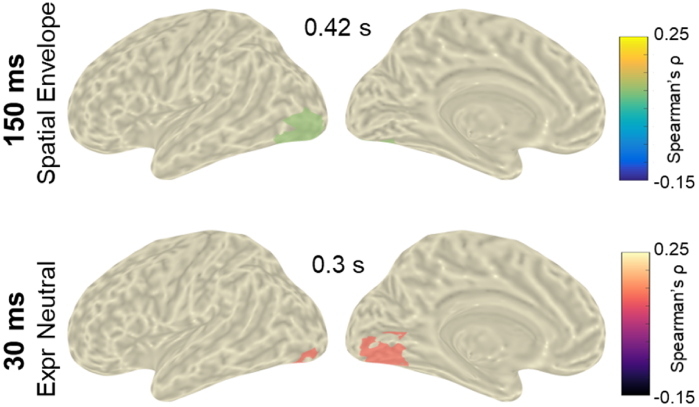
Significant correlations between: (1) MEG patterns for the 150 ms condition and the spatial envelope model RDM (**top**); (2) MEG patterns for the 30 ms condition and the high-level neutral-vs-others model (**bottom**). Only left hemisphere searchlights correlate with the two models. Maps are shown for the onset time of significant correlation, as clusters are sustained until offset (top: 0.54 s, bottom: 0.36 s).

The findings we present here constitute a stepping stone towards a better understanding of high-level representations in face perception. While binary categorical models can estimate high-level representations and task-related processing, the code supporting visual perception is likely to be better understood in terms of behavioural goals and the visual features supporting them. We show that face-responsive cortex dynamically encodes facial configuration starting with first-order features, and that this supports behavioural representations when participants are performing an expression discrimination task. Furthermore, we show that the cascade of processing stages changes with stimulus duration, pointing to the adaptability of the face processing system in achieving goals with limited visual input. This highlights the importance of investigating neural computations in a spatiotemporally resolved fashion; furthermore, when employing rapid presentation paradigms, it is important to consider the changes in neural dynamics and stimulus representations induced by relatively small changes in stimulus duration. Together, our results bridge findings from previous fMRI and electrophysiological research, revealing the spatiotemporal structure of face representations in human occipitotemporal cortex.

## Supporting information

SourceMovies1

## Acknowledgements

The authors would like to thank Lorenzo Magazzini and Gavin Perry for advice on the study design, and Dimitrios Pantazis for helpful comments on the manuscript. The study was supported by the UK MEG Partnership Grant (MRC/EPSRC, MR/K005464/1), CUBRIC and the School of Psychology at Cardiff University.

## Conflict of interest

The authors declare no competing interests.

## Appendix

**Supplementary Table 1:** Face decoding results.

|  | Sensor-space |  |  | Source-space |
| --- | --- | --- | --- | --- |
|  | 150 ms | 30 ms | 10 ms | 10 ms |
| Max % accuracy | 82.3 | 76.8 | 56.8 | 59.62 |
| SD (%) | 13.6 | 14.18 | 9.3 | 8.35 |
| Decoding onset (ms) | 100 | 100 | 147 | 120-150 |

**Supplementary Table 2:** Expression decoding results.

|  | Stimulus duration |  |  |  |  |  |  |  |  |  |  |  |
| --- | --- | --- | --- | --- | --- | --- | --- | --- | --- | --- | --- | --- |
|  | 150 ms |  |  | 30 ms |  |  | 10 ms |  |  | 30 + 150 ms |  |  |
|  | A-N | H-N | A-H | A-N | H-N | A-H | A-N | H-N | A-H | A-N | H-N | A-H |
| Max % accuracy | 61.9 | 63.1 | 60.76 | 57.79 | 58.49 | 58.12 | 56.62 | 55.86 | 55.87 | 60.48 | 60.21 | 59.74 |
| SD (%) | 8.57 | 6.78 | 9.34 | 10.91 | 9.92 | 10.38 | 10.88 | 9.11 | 13.66 | 9.04 | 10.52 | 13.41 |
| Decoding onset (ms) | 180 | 113 | 220 | 437 | 120 | 633 | N/A | N/A | N/A | 107 | 113 | 117 |
|  | Perceptual rating |  |  |  |  |  |  |  |  |  |  |  |
|  | 2 |  |  | 1 |  |  | 0 |  |  | 2 + 1 |  |  |
|  | A-N | H-N | A-H | A-N | H-N | A-H | A-N | H-N | A-H | A-N | H-N | A-H |
| Max % accuracy | 59.55 | 62.54 | 64.03 | 56.56 | 56.88 | 56.63 | 57.64 | 55.32 | 56.01 | 60.43 | 62.25 | 60.24 |
| SD (%) | 12.24 | 11.6 | 10.82 | 12.1 | 13.63 | 13.21 | 14.46 | 10.24 | 12.47 | 11.95 | 12.07 | 12.25 |
| Decoding onset (ms) | 230 | 113 | 523 | 307 | 120 | 130 | N/A | N/A | N/A | 220 | 113 | 127 |

**Supplementary Table 3:** RSA results for the 5 models achieving significant correlations.

| *Model* | Behavioural | Expression |  | Spatial Envelope |  | Global Config |  | Local Config |  |
| --- | --- | --- | --- | --- | --- | --- | --- | --- | --- |
| **150 ms** | $\rho$ | $\rho$ | $\rho_{part}$ | $\rho$ | $\rho_{part}$ | $\rho$ | $\rho_{part}$ | $\rho$ | $\rho_{part}$ |
| Max rho | 0.23 | 0.14 | 0.14 | 0.17 | 0.17 | 0.18 | 0.17 | 0.18 | 0.16 |
| SD | 0.12 | 0.03 | 0.03 | 0.06 | 0.06 | 0.09 | 0.08 | 0.07 | 0.06 |
| Onset (ms) | 120 | N/A | N/A | 420 | 390 | 300 | 330 | 120 | 120 |
| Offset (ms) | 390 | N/A | N/A | 540 | 540 | 450 | 390 | 300 | 180 |
| **30 ms** | $\rho$ | $\rho$ | $\rho_{part}$ | $\rho$ | $\rho_{part}$ | $\rho$ | $\rho_{part}$ | $\rho$ | $\rho_{part}$ |
| Max rho | 0.17 | 0.14 | 0.14 | 0.13 | 0.12 | 0.15 | 0.13 | 0.14 | 0.12 |
| SD | 0.07 | 0.03 | 0.03 | 0.05 | 0.04 | 0.07 | 0.03 | 0.05 | 0.04 |
| Onset (ms) | 210 | 300 | 300 | N/A | N/A | 390 | N/A | 360 | N/A |
| Offset (ms) | 540 | 360 | 360 | N/A | N/A | 450 | N/A | 510 | N/A |
| **10 ms** | $\rho$ | $\rho$ | $\rho_{part}$ | $\rho$ | $\rho_{part}$ | $\rho$ | $\rho_{part}$ | $\rho$ | $\rho_{part}$ |
| Max rho | 0.18 | 0.09 | 0.1 | 0.13 | 0.14 | 0.17 | 0.16 | 0.11 | 0.12 |
| SD | 0.04 | 0.03 | 0.03 | 0.04 | 0.05 | 0.04 | 0.04 | 0.03 | 0.04 |
| Onset (ms) | 120 | N/A | N/A | N/A | N/A | N/A | N/A | N/A | 180 |
| Offset (ms) | 660 | N/A | N/A | N/A | N/A | N/A | N/A | N/A | 240 |

**Supplementary Table 4:**
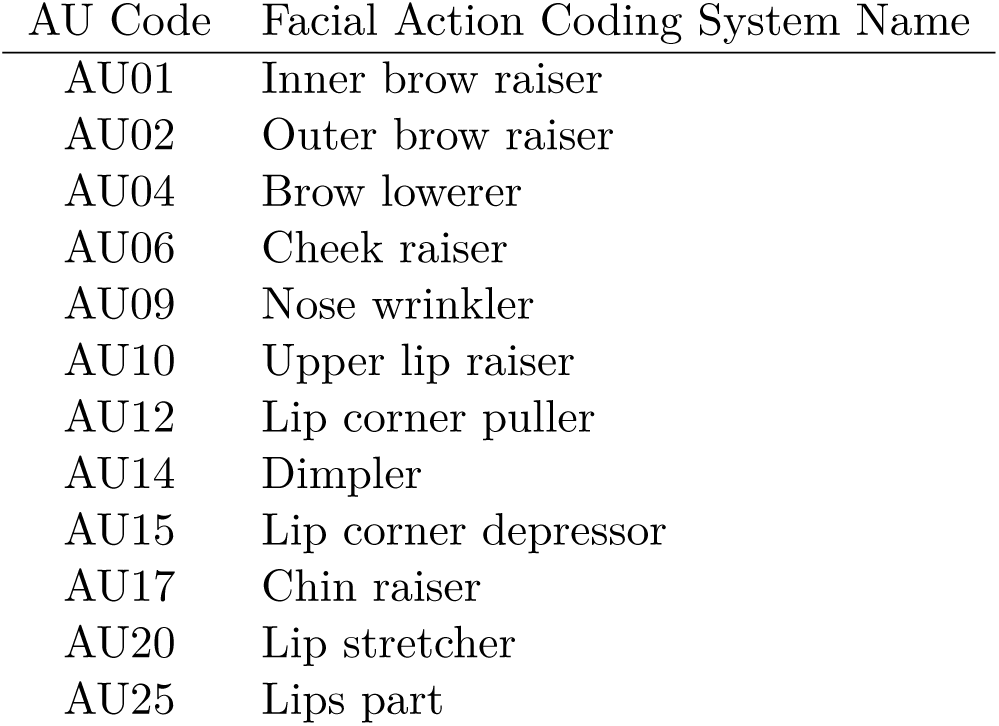
Action Units (AU) used to create the Action Unit model RDM.

## Supplementary Analysis 1: Event-related field (ERF) analysis

We assessed the presence of difference between conditions in event-related fields (ERF). For the purposes of this analysis, MEG data were bandpass-filtered between 0.1 and 30 Hz and axial gradiometer event-related fields were averaged across subjects to calculate the global field power across all trials and conditions. This allowed us to determine three time windows of interest for evoked response component analysis: 63-137 ms (M100), 137-203 ms (M170), and 203–306 ms (M220).

Next, we averaged evoked response fields for each condition and subject within the three time windows. We tested for differences between responses to faces and scrambled stimuli, and between responses to different emotional expressions, using paired t-tests and repeated-measures ANOVAs respectively at each sensor and time window. Significant sensors were determined using randomization testing (5000 iterations) and corrected for multiple comparisons using the maximal statistic distribution (*α* = 0.001).

We assessed the presence of a response to faces by contrasting neutral faces with scrambled stimuli at each stimulus duration. For 150 ms faces, we found significant differences at M170 latencies and M220 latencies (*P* < 0.0007, *t*(24) *>* 6.07), but no significant effects at M100 latencies surviving our alpha of 0.001 (only one occipital sen sor showed a non-significant effect with *P* = 0.0059, *t*(24) = 4.89). A significant, but smaller, cluster of right temporal sensors was also found for 30 ms faces at M170 latencies (*P* < 0.0004, *t*(24) *>* 5.99). No conclusive effects were found when contrasting faces presented for 10 ms with their scrambled counterparts, regardless of whether trials where a face was perceived were excluded or not (*P >* 0.015, *t*(24) *<* 4.66 across comparisons), and no effect of emotional expression was found at any of the stimulus durations (*P >* 0.06, *F* (2, 48) *<* 8.59). Several factors could explain the absence of emotional expression effects in our ERF data: (1) stimuli were highly controlled for low-level properties, minimizing visually-driven differences in early time windows; (2) our time windows of interest did not include late stages dominated by task-related processing of expression; (3) we performed a whole-brain analysis with a conservative correction for multiple comparisons.

**Supplementary Figure 1:**
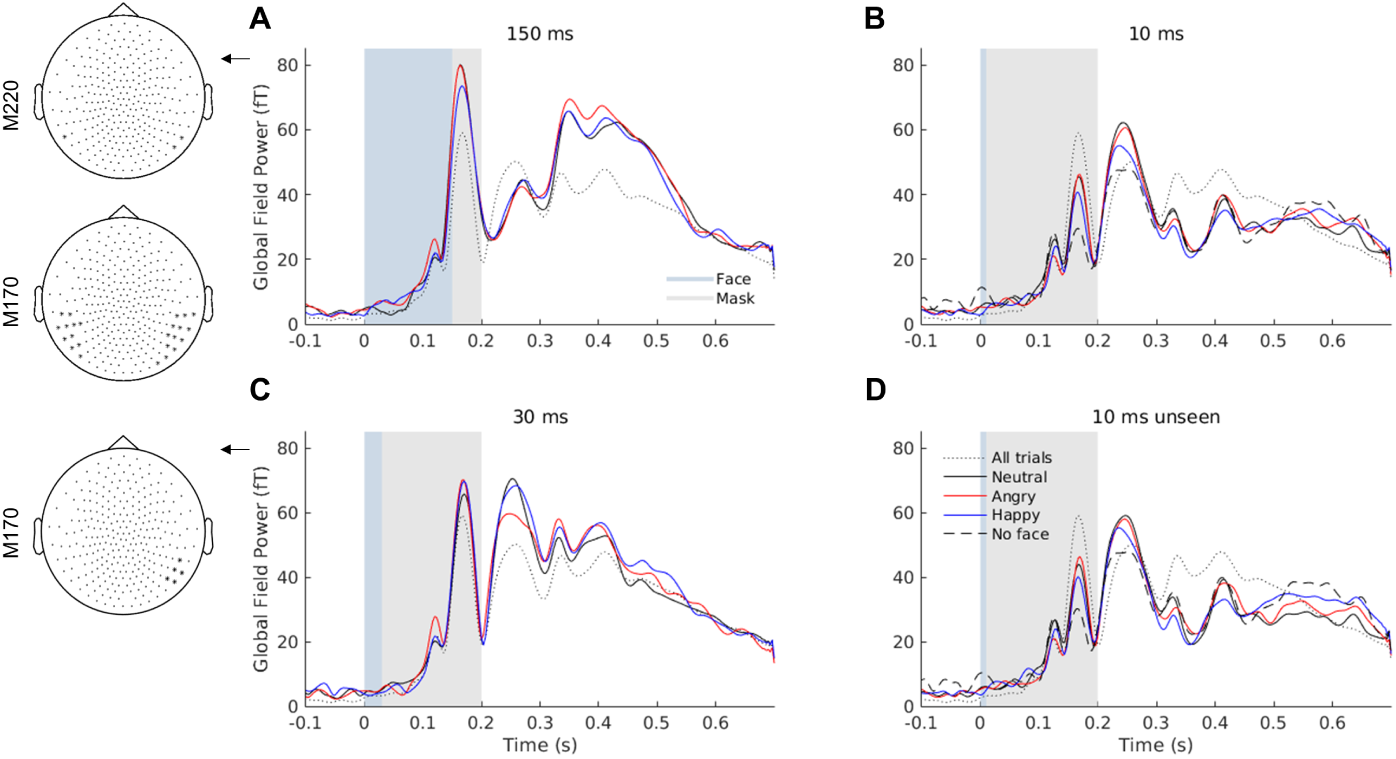
ERF analysis results. **A-D.** Global field power averaged across participants and trials for each stimulus duration condition. Note decreasing M170 amplitudes with stimulus duration. **Left.** Significant sensors in the face vs scrambled contrast at M170 (137-203 ms) and M220 (203-306 ms) latencies (*P* < 0.001 corrected).

**Supplementary Figure 2:**
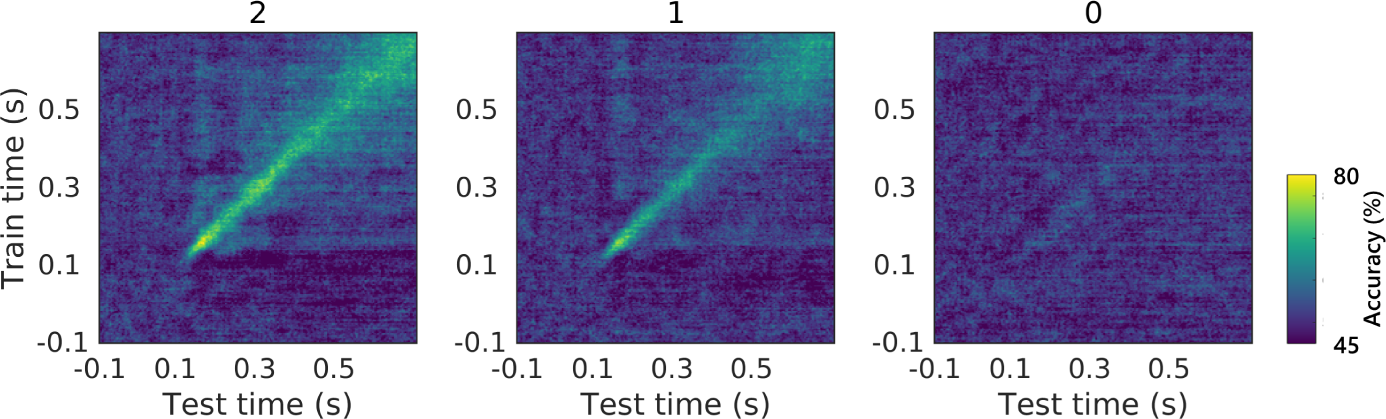
Face vs scrambled temporal generalization decoding for each perceptual rating category. The same progression from stable to transient representations is observed as when datasets were split according to stimulus duration.

**Supplementary Figure 3:**
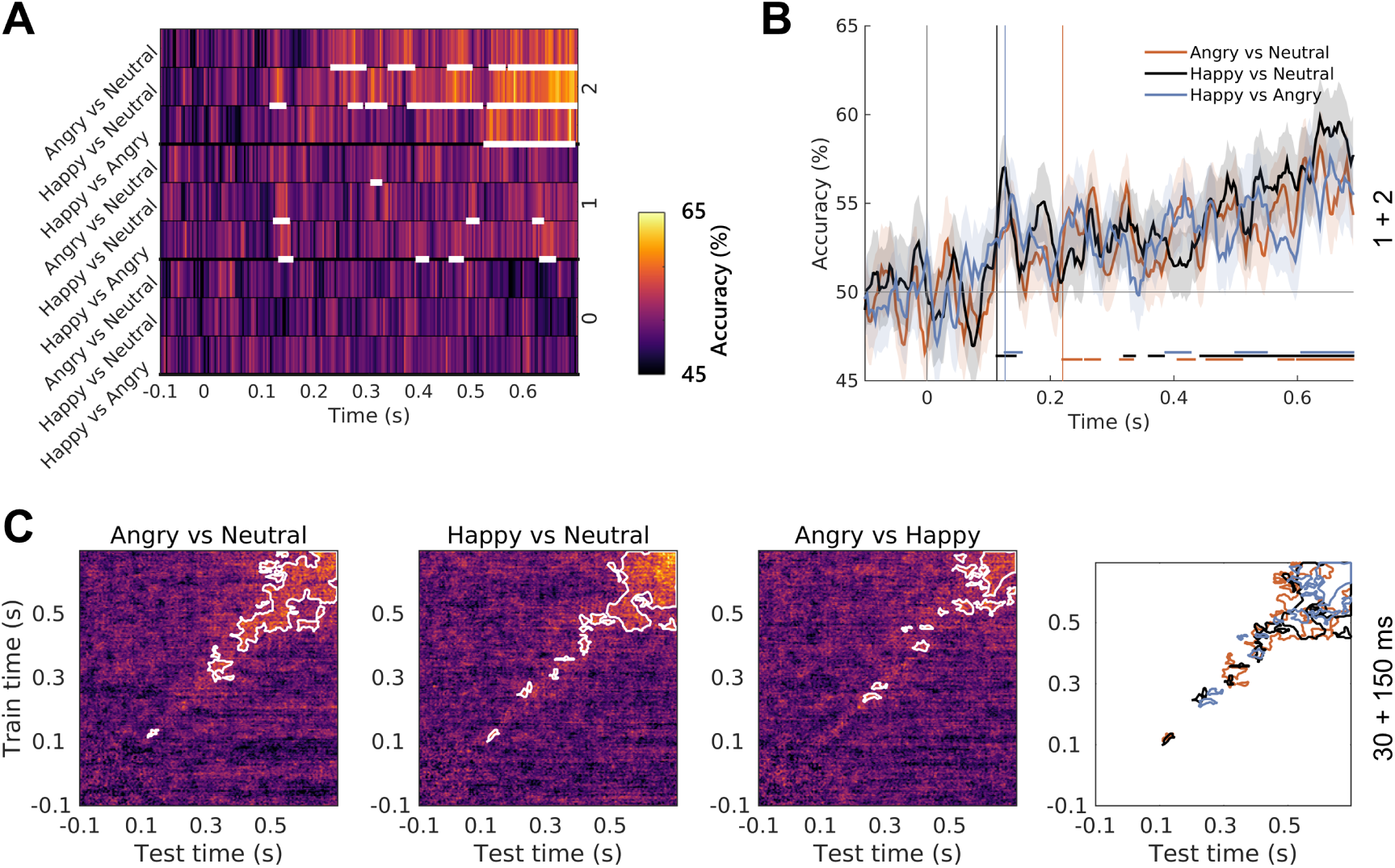
Expression decoding. **A.** Time-resolved decoding accuracy for each pair of expressions and perceptual rating, with above-chance time-windows highlighted in white (*P* < 0.05 corrected). **B.** Accuracy time-courses obtained using pooled datasets (awareness ratings of 1 + 2). **C.** Temporal generalization accuracy and signifi cant clusters (white contours; *P* < 0.05, corrected) for the three decoding problems using the pooled datasets (duration of 30 + 150 ms). The last panel shows significant temporal generalization clusters for all three decoding problems. Angry vs neutral decoding leads to earlier stable representations.

**Supplementary Figure 4:**
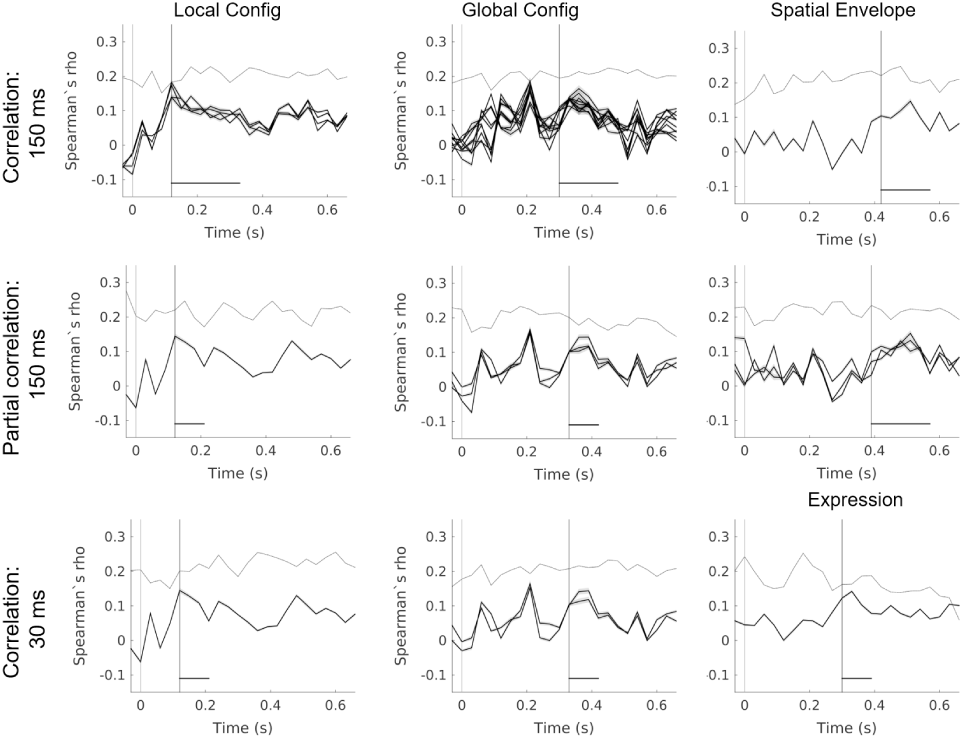
Correlation time-courses obtained in the RSA analysis. All significant searchlights are plotted separately against a noise ceiling averaged across significant searchlights.

## References

Aguado, L., Valdés-Conroy, B., Rodríguez, S., Román, F. J., Diéguez-Risco, T., & Fernández-Cahill, M. (2012). Modulation of early perceptual processing by emotional expression and acquired valence of faces: An ERP study. Journal of Psychophysiology, 26 (1), 29–41. doi:10.1027/0269-8803/a000065

Axelrod, V. (2010). The Fusiform Face Area: In Quest of Holistic Face Processing. Journal of Neuroscience, 30 (26), 8699–8701. doi:10.1523/JNEUROSCI.1921-10.2010

Axelrod, V., Bar, M., & Rees, G. (2015). Exploring the unconscious using faces. Trends in Cognitive Sciences, 19 (1), 35–45. doi:10.1016/j.tics.2014.11.003

Baltrusaitis, T., Robinson, P., & Morency, L.-P. (2016). OpenFace: an open source facial behaviour analysis toolkit. 2016 IEEE Winter Conference on Applications of Computer Vision (WACV), 1–10.

Bankson, B. B., Hebart, M. N., Groen, I. I., & Baker, C. I. (2018). The temporal evolution of conceptual object representations revealed through models of behavior, semantics and deep neural networks. NeuroImage, 178, 172–182. doi:10.1016/J.NEUROIMAGE.2018.05.037

Boehler, C. N., Schoenfeld, M. A., Heinze, H.-J., & Hopf, J.-M. (2008). Rapid recurrent processing gates awareness in primary visual cortex. Proceedings of the National Academy of Sciences, 105 (25), 8742–8747.

Calder, A. J., Young, A. W., Keane, J., & Dean, M. (2000). Configural information in facial expression perception. Journal of Experimental Psychology: Human Perception and Performance, 26 (2), 527–551. doi:10.1037/0096-1523.26.2.527

Chang, L. & Tsao, D. Y. (2017). The Code for Facial Identity in the Primate Brain. Cell, 169 (6), 1013–1028. doi:10.1016/j.cell.2017.05.011

Dehaene, S. (2016). Decoding the Dynamics of Conscious Perception : The Temporal Generalization Method. In B. G & C. Y (Eds.), Micro-, meso- and macro-dynamics of the brain (pp. 85–97). New York: Springer. doi:10.1007/978-3-319-28802-4

Diamond, R. & Carey, S. (1986). Why faces are and are not special: an effect of expertise. Journal of experimental psychology, 115 (2), 107–117.

Dima, D. C., Perry, G., Messaritaki, E., Zhang, J., & Singh, K. D. (2018). Spatiotemporal dynamics in human visual cortex rapidly encode the emotional content of faces. Human Brain Mapping, 39 (10), 3993–4006. doi:10.1002/hbm.24226

Dima, D. C., Perry, G., & Singh, K. D. (2018). Spatial frequency supports the emergence of categorical representations in visual cortex during natural scene perception. NeuroImage, 179, 102–116. doi:10.1016/j.neuroimage.2018.06.033

Eimer, M., Gosling, A., Nicholas, S., & Kiss, M. (2011). The N170 component and its links to configural face processing: A rapid neural adaptation study. Brain Research, 1376, 76–87. doi:10.1016/J.BRAINRES.2010.12.046

Ekman, P. & Friesen, W. (1977). Facial action coding system: a technique for the measurement of facial movement. Palo Alto, CA: Consulting Psychologists Press.

Fan, R.-E., Chang, K.-W., Hsieh, C.-J., Wang, X.-R., & Lin, C.-J. (2008). LIBLINEAR: A Library for Large Linear Classification. Journal of Machine Learning Research, 9 (2008), 1871–1874. doi:10.1038/oby.2011.351

Farah, M. J., Wilson, K. D., & Tanaka, J. N. (1998). What Is “ Special “ About Face Perception ? Psychological review, 105 (3), 482–498. doi:10.1037//0033-295X.105.3.482

Freiwald, W., Duchaine, B., & Yovel, G. (2016). Face Processing Systems: From Neurons to Real-World Social Perception. Annual Review of Neuroscience, 39 (1), 325–346. doi:10.1146/annurev-neuro-070815-013934

Furl, N., Lohse, M., & Pizzorni-Ferrarese, F. (2017). Low-frequency oscillations employ a general coding of the spatio-temporal similarity of dynamic faces. NeuroImage, 157, 486–499. doi:10.1016/j.neuroimage.2017.06.023

Gohel, B., Lim, S., Kim, M.-Y., Kwon, H., & Kim, K. (2018). Dynamic pattern decoding of source-reconstructed MEG or EEG data: Perspective of multivariate pattern analysis and signal leakage. Computers in Biology and Medicine, 93, 106–116. doi:10.1016/j. compbiomed.2017.12.020

Golarai, G., Ghahremani, D. G., Eberhardt, J. L., & Gabrieli, J. D. E. (2015). Distinct representations of configural and part information across multiple face-selective regions of the human brain. Frontiers in Psychology, 6, 1710. doi:10.3389/fpsyg.2015.01710

Greene, M. R., Baldassano, C., Esteva, A., Beck, D. M., & Fei-fei, L. (2016). Visual Scenes are Categorized by Function. Journal of Experimental Psychology: General, 145 (1), 82–94. doi:10.1037/xge0000129.Visual

Grill-Spector, K., Weiner, K. S., Gomez, J., Stigliani, A., & Natu, V. S. (2018). The functional neuroanatomy of face perception: From brain measurements to deep neural networks. Interface Focus, 8. doi:10.1098/rsfs.2018.0013

Groen, I. I., Greene, M. R., Baldassano, C., Fei-Fei, L., Beck, D. M., & Baker, C. I. (2018). Distinct contributions of functional and deep neural network features to representational similarity of scenes in human brain and behavior. eLife, 7, e32962. doi:10.7554/eLife.32962

Grootswagers, T., Wardle, S. G., & Carlson, T. A. (2017). Decoding Dynamic Brain Patterns from Evoked Responses: A Tutorial on Multivariate Pattern Analysis Applied to Time Series Neuroimaging Data. Journal of Cognitive Neuroscience, 29 (4), 677– 697. doi:10.1162/jocn{\_}a{\_}01068

Guggenmos, M., Sterzer, P., & Cichy, R. M. (2018). Multivariate pattern analysis for MEG: A comparison of dissimilarity measures. NeuroImage, 173, 434–447. doi:10. 1016/J.NEUROIMAGE.2018.02.044

Harris, A. M. & Aguirre, G. K. (2008). The effects of parts, wholes, and familiarity on face-selective responses in MEG. Journal of Vision, 8 (10), 4–4. doi:10.1167/8.10.4

Hedger, N., Gray, K. L. H., Garner, M., & Adams, W. J. (2016). Are visual threats prioritised without awareness? A critical review and meta analysis involving 3 behavioural paradigms and 2696 observers. Psychological Bulletin, 142 (9), 934–968.

Henriksson, L., Mur, M., & Kriegeskorte, N. (2015). Faciotopy-A face-feature map with face-like topology in the human occipital face area. Cortex, 72, 156–167. doi:10.1016/ j.cortex.2015.06.030

Hillebrand, A., Barnes, G. R., Bosboom, J. L., Berendse, H. W., & Stam, C. J. (2012). Frequency-dependent functional connectivity within resting-state networks : An atlas-based MEG beamformer solution. NeuroImage, 59 (4), 3909–3921. doi:10.1016/j.neuroimage.2011.11.005

Hillebrand, A., Singh, K. D., Holliday, I. E., Furlong, P. L., & Barnes, G. R. (2005). A new approach to neuroimaging with magnetoencephalography. Human Brain Mapping, 25 (2), 199–211. doi:10.1002/hbm.20102

King, J.-R. & Dehaene, S. (2014). Characterizing the dynamics of mental representations : the temporal generalization method. Trends in Cognitive Sciences, 18 (4), 203–210. doi:10.1016/j.tics.2014.01.002

Koster, E. H. W., Verschuere, B., Burssens, B., Custers, R., & Crombez, G. (2007). Attention for Emotional Faces Under Restricted Awareness Revisited : Do Emotional Faces Automatically Attract Attention ? Emotion, 7 (2), 285–295. doi:10.1037/1528-3542.7.2.285

Lamme, V. A. F., Zipser, K., & Spekreijse, H. (2002). Masking Interrupts Figure-Ground Signals in V1. Journal of Cognitive Neuroscience, 14 (7), 1044–1053. doi:10.1162/089892902320474490

Leopold, D. A., O’Toole, A. J., Vetter, T., & Blanz, V. (2001). Prototype-referenced shape encoding revealed by high-level aftereffects. Nature Neuroscience, 4 (1), 89–94. doi:10.1038/82947

Longmore, C. A., Liu, C. H., & Young, A. W. (2015). The importance of internal facial features in learning new faces. Quarterly Journal of Experimental Psychology, 68 (2), 249–260. doi:10.1080/17470218.2014.939666

Maurer, D., Grand, R. L., & Mondloch, C. J. (2002). The many faces of configural processing. Trends in Cognitive Sciences, 6 (6), 255–260. doi:10.1016/S1364-6613(02)01903-4

Mercure, E., Dick, F., & Johnson, M. H. (2008). Featural and configural face processing differentially modulate ERP components. Brain Research, 1239, 162–170. doi:10.1016/J.BRAINRES.2008.07.098

Micallef, L. & Rodgers, P. (2014). eulerAPE: Drawing Area-Proportional 3-Venn Diagrams Using Ellipses. PLoS ONE, 9 (7), e101717. doi:10.1371/journal.pone.0101717

Mohsenzadeh, Y., Qin, S., Cichy, R. M., & Pantazis, D. (2018). Ultra-Rapid serial visual presentation reveals dynamics of feedforward and feedback processes in the ventral visual pathway. eLife, 7 (e36329). doi:10.7554/eLife.36329.001

Nichols, T. E. & Holmes, A. P. (2001). Nonparametric Permutation Tests For Functional Neuroimaging : A Primer with Examples. Human Brain Mapping, 25 (15), 1–25. doi:10.1002/hbm.1058

Nili, H., Wingfield, C., Walther, A., Su, L., Marslen-Wilson, W., & Kriegeskorte, N. (2014). A Toolbox for Representational Similarity Analysis. PLoS Computational Biology, 10 (4), e1003553. doi:10.1371/journal.pcbi.1003553

Oliva, A. & Torralba, A. (2001). Modeling the Shape of the Scene : A Holistic Representation of the Spatial Envelope. International Journal of Computer Vision, 42 (3), 145–175. doi:10.1023/A:1011139631724

Oostenveld, R., Fries, P., Maris, E., & Schoffelen, J. M. (2011). FieldTrip: Open source software for advanced analysis of MEG, EEG, and invasive electrophysiological data. Computational Intelligence and Neuroscience, 2011, 156869. doi:10.1155/2011 /156869

Perry, G. & Singh, K. D. (2014). Localizing evoked and induced responses to faces using magnetoencephalography. European Journal of Neuroscience, 39 (9), 1517–1527. doi:10.1111/ejn.12520

Pessoa, L., Japee, S., & Sturman, D. (2006). Target Visibility and Visual Awareness Modulate Amygdala Responses to Fearful Faces. Cerebral Cortex, 16 (March), 366–375. doi:10.1093/cercor/bhi115

Piepers, D. W. & Robbins, R. A. (2012). A review and clarification of the terms “holistic,” “configural,” and “relational” in the face perception literature. Frontiers in Psychology, 3, 1–11. doi:10.3389/fpsyg.2012.00559

Said, C. P., Moore, C. D., Engell, A. D., & Haxby, J. V. (2018). Distributed representations of dynamic facial expressions in the superior temporal sulcus. Journal of Vision, 10 (5), 1–12. doi:10.1167/10.5.11.Introduction

Sato, M., Yamashita, O., Sato, M.-a., & Miyawaki, Y. (2018). Information spreading by a combination of MEG source estimation and multivariate pattern classification. PLOS ONE, 13 (6), e0198806. doi:10.1371/journal.pone.0198806

Schlossmacher, I., Junghöfer, M., Straube, T., & Bruchmann, M. (2017). No differential effects to facial expressions under continuous flash suppression: An event-related potentials study. NeuroImage, 163, 276–285. doi:10.1016/j.neuroimage.2017.09.034

Singh, K. D., Barnes, G. R., & Hillebrand, A. (2003). Group imaging of task-related changes in cortical synchronisation using nonparametric permutation testing. NeuroImage, 19, 1589–1601. doi:10.1016/S1053-8119(03)00249-0

Stelzer, J., Chen, Y., & Turner, R. (2013). Statistical inference and multiple testing correction in classification-based multi-voxel pattern analysis (MVPA): Random permutations and cluster size control. NeuroImage, 65, 69–82. doi:10.1016/j.neuroimage. 2012.09.063

Studebaker, G. (1985). A “rationalized” arcsine transform. Journal of speech and hearing research, 28, 455–462. doi:10.1044/jshr.2803.455

Su, L., Fonteneau, E., Marslen-wilson, W., & Kriegeskorte, N. (2012). Spatiotemporal Searchlight Representational Similarity Analysis in EMEG Source Space. In Second international workshop on pattern recognition in neuroimaging spatiotemporal. doi:10.1109/PRNI.2012.26

Tottenham, N., Tanaka, J. W., Leon, A. C., McCarry, T., Nurse, M., Hare, T. a., … Nelson, C. (2009). The NimStim set of facial expressions: judgments from untrained research participants. Psychiatry research, 168 (3), 242–9. doi:10.1016/j.psychres.2008.05.006

Tzourio-Mazoyer, N., Landeau, B., Papathanassiou, D., Crivello, F., Etard, O., Delcroix, N., … Joliot, M. (2002). Automated Anatomical Labeling of Activations in SPM Using a Macroscopic Anatomical Parcellation of the MNI MRI Single-Subject Brain. NeuroImage, 15, 273–289. doi:10.1006/nimg.2001.0978

Van Veen, B., van Drongelen, W., Yuchtman, M., & Suzuki, A. (1997). Localization of brain electrical activity via linearly constrained minimum variance spatial filtering. IEEE Transactions on Biomedical engineering, 44 (9), 867–880. doi:10.1109/10.623056

Vidaurre, D., Myers, N., Stokes, M., Nobre, A. C., & Woolrich, M. W. (2018). Temporally unconstrained decoding reveals consistent but time-varying stages of stimulus processing. bioRxiv, 260943. doi:10.1101/260943

Visconti Di Oleggio Castello, M., Wheeler, K. G., Cipolli, C., & Gobbini, I. (2017). Familiarity facilitates feature-based face processing. PLoS One, 12 (6), e0178895. doi:10. 1371/journal.pone.0178895

Vrba, J. & Robinson, S. E. (2001). Signal processing in magnetoencephalography. Methods, 25 (2), 249–271. doi:10.1006/meth.2001.1238

Walther, A., Nili, H., Ejaz, N., Alink, A., Kriegeskorte, N., & Diedrichsen, J. (2016). Reliability of dissimilarity measures for multi-voxel pattern analysis. NeuroImage, 137, 188–200. doi:10.1016/j.neuroimage.2015.12.012

Walther, D. B., Caddigan, E., Fei-Fei, L., & Beck, D. M. (2009). Natural Scene Categories Revealed in Distributed Patterns of Activity in the Human Brain. Journal of Neuroscience, 29 (34), 10573–10581. doi:10.1523/JNEUROSCI.0559-09.2009

Wegrzyn, M., Vogt, M., Kireclioglu, B., Schneider, J., & Kissler, J. (2017). Mapping the emotional face. How individual face parts contribute to successful emotion recognition. PLoS ONE, 12 (5), 1–15.

Willenbockel, V., Sadr, J., Fiset, D., Horne, G. O., Gosselin, F., & Tanaka, J. W. (2010). Controlling low-level image properties: The SHINE toolbox. Behavior Research Methods, 42 (3), 671–684. doi:10.3758/BRM.42.3.671

Wyatte, D., Jilk, D. J., & O’Reilly, R. C. (2014). Early recurrent feedback facilitates visual object recognition under challenging conditions. Frontiers in Psychology, 5, 674. doi:10.3389/fpsyg.2014.00674

Zachariou, V., Safiullah, Z. N., & Ungerleider, L. G. (2018). The Fusiform and Occipital Face Areas Can Process a Nonface Category Equivalently to Faces. Journal of Cognitive Neuroscience, 30 (10), 1499–1516. doi:10.1162/jocn{\_}a{\_}01288

Zion-Golumbic, E. & Bentin, S. (2007). Dissociated Neural Mechanisms for Face Detection and Configural Encoding: Evidence from N170 and Induced Gamma-Band Oscillation Effects. Cerebral Cortex, 17 (8), 1741–1749. doi:10.1093/cercor/bhl100

